# Comprehensive assessment of Indian variations in the druggable kinome landscape highlights distinct insights at the sequence, structure and pharmacogenomic stratum

**DOI:** 10.1101/2021.05.23.445314

**Authors:** Gayatri Panda, Neha Mishra, Disha Sharma, Rintu Kutum, Rahul C. Bhoyar, Abhinav Jain, Mohamed Imran, Vigneshwar Senthilvel, Mohit Kumar Divakar, Anushree Mishra, Parth Garg, Priyanka Banerjee, Sridhar Sivasubbu, Vinod Scaria, Arjun Ray

**Affiliations:** Department of Computational Biology, Indraprastha Institute of Information Technology, Okhla, India; Academy of Scientific and Innovative Research (AcSIR), Ghaziabad, India; CSIR-Institute of Genomics and Integrative Biology, Mathura Road, Delhi-110020, India; Institute for Physiology, Charité-University Medicine Berlin, 10115 Berlin, Germany

**Keywords:** Indian genetic variations, IndiGenome Consortium, Pharmacogenomics, single nucleotide varinants, docking, adverse drug reactions

## Abstract

India confines more than 17% of the world’s population and has a diverse genetic makeup with several clinically relevant rare mutations belonging to many sub-group which are undervalued in global sequencing datasets like the 1000 Genome data (1KG) containing limited samples for Indian ethnicity. Such databases are critical for the pharmaceutical and drug development industry where the diversity plays a crucial role in identifying genetic disposition towards adverse drug reactions. A qualitative and comparative sequence and structural study utilizing variant information present in the recently published, largest curated Indian genome database (IndiGen) and the 1000 Genome data was performed for variants belonging to the kinase coding genes,the second most targeted group of drug targets. The sequence level analysis identified similarities and differences among different populations based on the nsSNVs and amino acid exchange frequencies whereas comparative structural analysis of IndiGen variants was performed with pathogenic variants reported in UniProtKB Humsavar data. The influence of these variations on structural features of the protein, such as structural stability, solvent accessibility, hydrophobicity, and the hydrogen-bond network were investigated. In-silico screening of the known drugs to these Indian variation-containing proteins reveal critical differences imparted in the strength of binding due to the variations present in the Indian population. In conclusion, this study constitutes a comprehensive investigation into the understanding of common variations present in the second largest population in the world, and investigating its implications in the sequence, structural and pharmacogenomic landscape. The preliminary investigation reported in this paper, supporting the screening and detection of ADRs specific to the Indian population could aid in the development of techniques for pre-clinical and post-market screening of drug-related adverse events in the Indian population.

## Introduction

The presence of single nucleotide polymorphisms imparts a genetic basis of human complex diseases and human phenotypic variations [Marian, 2013]. Those single nucleotide variants (SNVs) which are present in more that 1 % of the population qualify as a single nucleotide polymorphisms (SNPs). As per various reports, SNPs are found to be responsible for defining the risk of an individual’s susceptibility to various drug responses and illnesses [Alwi, 2005]. The distribution of allele frequency of SNVs provides relevant information about the evolution, migration, and genetic structure of a population [Bomba et al., 2017, Sanghera et al., 2008]. Genetic variability is among many factors contributing to the inter-individual differences in drugresponse and individuals of various geographic ancestry exhibit genetic variations with varying frequencies [Henn et al., 2016, Lauschke et al., 2019, Schärfe et al., 2017]. Thus, an individual’s risk differs for many drugs with respect to geographic ancestry. Those variants which have a functional effect in a target for commonly prescribed drugs could alter drug pharmacokinetics (PK) and pharmacodynamics (PD) resulting in adverse-drug reaction. Most of the genetic variant-related data come from databases like the 1000 Genome database [Auton et al., 2015] and gnomAD [Karczewski et al., 2020] database containing ethnicity-wise variant information which is largely Eurocentric. It is so because majority of the studies that are performed to associate genetic variants with diseases, like the Genome-Wide Association Studies (GWAS) have been conducted mainly on the European population (78%) followed by Asian(10%), African(2%), Hispanic(1%), and other ethnicities (<1%) [Sirugo et al., 2019] neglecting the Indian population. It creates an information bias leading to a population-specific disease assessment analysis leaving the African and Indian populations under-studied and under-consulted. These population-specific SNVs deviate in variation patterns from other over-represented populations causing health and diagnosis disparities[Chan et al., 2015] [Wei et al., 2012].

Globally, Adverse drug reactions (ADRs) are a major contributor to morbidity and mortality [Khalil and Huang, 2020]. The presence of a genomic variation in genes coding for drug transport and metabolism have been associated with inter-individual differences in drug response and ADR risks. Several SNV-related studies have shown that variants can modulate the efficacy of a drug leading to adverse drug reactions (ADRs) [Impicciatore et al., 2001] [Sanghera et al., 2008]. Drug Gene Interaction Database (DGIdb) organizes the drug-gene interactions from various papers, databases and web resources[Freshour et al., 2021]. dbSNP [Sherry et al., 2001a], a curated database alone contains 38 million SNPs which makes timely maintenance, integration, and correction a cumbersome process [Sherry et al., 2001b]. SNVs are a vital and decisive factor for finalizing a therapeutic approach and selection of drug and their dosages [Alwi, 2005]. European population being primary conduct of drug trials prior to approval and marketing of drugs could be one of the factors on the occurrence of ADRs[Clinical and Guidelines, 2006]. Hence, this prioritizes the need for population-specific pharmacogenomic analysis and integration of gene, drug, pathway, and potential drug-target related information. Genetic studies of populations from the Indian subcontinent are important due to India’s large share of the global population, complex demographic background, and unique social structure. Indo-genomic variation is fascinating due to the diverse ancestral components, social categorization of people, endogamy practised in different cultures, and dynamic and ancient admixture events that the Indian population has experienced over a long period of time.[Bamshad et al., 2001]. Reports suggest that the population expansion in India (post-agriculture) has led to the emergence of a huge amount of genomic diversity exceeding the genetic diversity of the whole of Europe[Sengupta et al., 2016]

The practice of endogamy in various communities disturbs the frequency of a disease in different sub-groups of the Indian population [Nakatsuka et al., 2017], indicating that genetic divergence can also affect the efficacy of the drug. Globally, India is the largest generic drug provider [Bhosle et al., 2016]. Regardless of the Indian genetic diversity, the current healthcare system in India follows the same drug therapy as in Europe and America. The use of genetic information, experiments, and other types of molecular screening helps a practitioner to choose an appropriate therapy for the first time, avoiding the time-consuming and expensive trial-and-error medication cycle. Extensive research on the population diversities and related SNVs causing the different inter-individual drug responses is the need of the hour for efficient treatment design. IndiGen programme was initiated with an aim to collect sequencing data of thousands of individuals from diverse ethnic groups in India and develop public health technologies applications using this population genome data[Jain et al., 2021].

In our present work, we conducted an exhaustive and comparative study of common Indian-specific variants (using IndiGen data) with other populations to identify the population-specific variations causing a difference in drug responses and ADRs. This pharmacogenomic study was executed by keeping a focus on druggable genes of kinase’s family, the second most targeted group of drug targets after the G-protein coupled receptors [Bhullar et al., 2018]. The human genome encodes 538 protein kinases[Bhullar et al., 2018]. Many of these kinases are associated with deadly diseases like cancer [Paul and Mukhopadhyay, 2012]. Most of the kinase targeting drugs have been tested and approved based on the trials done on European populations and it is possible that the same drugs might exhibit a deviation in efficacy and response on Indian population. The presence of a SNV in functionally important genes have higher chances of deleterious impact by either affecting drug-gene interaction or by causing structural changes at the protein level leading to disruption of the drug-binding sites [Lee, 2010]. As a result, interpreting the number of mutations and their effect on the structure, stability, and function of the protein is crucial. Any destabilising non-synonymous SNV (nsSNV) will cause the drug’s metabolic process to be disrupted. This study was carried out at both sequence and structure level to examine the effect of missense mutations in Drug-Gene interaction as well as the structural changes caused by these mutations at the protein level.The sequence-level analysis was implemented to perceive the similarities and differences among different populations based on the single nucleotide variants (SNVs) and amino acid exchange frequencies. The effect of these variants on structural properties of the protein, like structural stability, solvent-accessibility, hydrophobicity, and the hydrogen-bond network were measured by utilizing different structural analysis tools. Any modification in protein-ligand binding due to the presence of SNVs was analyzed by molecular docking method. A comparative structural analysis was conducted using UniProtKB Humsavar data.This work will help us understand the variability caused by these variants and thus could guide us in deciphering the effect of SNV in the efficacy of the drug-protein/gene interaction.

## Materials and Methods

### Variant Data collection

The genetic variants and their allele frequencies in the Indian population were curated from 1029 whole-genome sequences of unrelated individuals across India collected as part of the IndiGen programme to represent diverse Indo-ethnicities [Jain et al., 2020]. The variant data consisted of single nucleotide variants and indels, which were annotated according to the GRCh38 human reference genome using Annovar[Wang et al., 2010]. Only non-synonymous variants (nsSNVs) were considered for our study [Jain et al., 2020]. The publicly available variant calling format (vcf) file 1000g2015aug all was downloaded for 1000 Genome data [Clarke et al., 2016].

### Assembling druggable genes

The Drug Gene Interaction Database (DGIdb) version 3 contains information on all currently approved drugs as well as other future targets of interest.[Freshour et al., 2021]. Genes were annotated in this database with respect to known drug-gene interactions and potential druggability. It normalizes its content from 30 open-source databases like DrugBank [Wishart et al., 2008], therapeutic target database (TTD)[Chen et al., 2002], PharmGKB [Boom et al., 2013] and other web resources like Oncology Knowledge Base (OncoKB) [With et al., 2017], cancer genome interpreter (CGI) [Tamborero et al., 2018], etc. A list of 545 druggable kinases and associated FDA-approved drugs were retrieved from the DGIdb using browse category search while limiting the categories to specific resources i.e ‘GuideToPharmacologyGenes’(Supplemental Table S1). The Guide to Pharmacology is a curated repository of ligand-activity-target relationships, with most of its information derived from high-quality pharmacological and medicinal literature. This druggable kinase gene list was further enriched by adding features like Ensembl ID, PDB ID, RefSeq Match Transcript, gene start - gene end, Uniprot ID, sequence length, and structure length, etc. using BioMart resource [Smedley et al., 2009]

### Data Preparation

#### Sequence Data Preparation

The dataset used for sequence analysis contained 545 druggable kinase genes (Supplemental Table S1) and their associated variants. Protein sequences for these genes were downloaded from NCBI Genbank, and mutant sequences were prepared by altering the native sequence according to the Annovar data.

#### Structure Data Preparation

The data for structural analysis was prepared by applying a few filters to the base sequence data. These filters were 1. Availability of protein crystal structure, 2. Availability of drug molecules against the protein, 3. Crystal structure and sequence coverage ≥ 70%, 4. Allele frequency of the nsSNV observed in the IndiGen population ≥ 10%, 5. SNV coverage to the crystal structure. After applying these filters, 12 genes and their corresponding 22 variants were left and were referred to as IndiGen Structure data (Supplemental Table S4). In an attempt to conduct a comparative structural analysis, Humsavar (Human polymorphisms and disease mutations) data was taken. It lists all missense variants annotated in human UniProtKB/Swiss-Prot entries (Release: 2020 04 of 12-Aug-2020). In this data, the variants were classified as disease-causing (31132-64.1%), Polymorphisms (39464-23%), and Unclassified (8381-12.9%). The variants associated with the genes present in IndiGen Structure data were extracted from Humsavar’s complete list of variants. This dataset was referred to as the Humsavar dataset, which consisted of 217 variants and was used for benchmarking structural analysis (Supplemental Table S13).

### Data Processing and Visualization

#### Drug, Gene and Variant Tree

The primary goal of this analysis was to have a quantitative and qualitative insight into the frequency of occurrence of variation in the family of kinases and their drug availability. This study will aid in gathering information related to the family of kinases with more variations and drugs reported. An online tool, KinMap [Eid et al., 2017], was used for an interactive exploration of kinase coding genes present in IndiGen data. The genes associated with 545 druggable kinases, the number of variations, and drugs reported against each gene in DGIdb were given as an input to this tool (Supplemental_ Table _S11).

#### Amino acid Conversions and Mutabilities

The tendency to convert an amino acid type to another type and identify any pattern in this conversion can helpto understand the change in a protein sequence’s physicochemical property. This analysis was conducted using a python script, and the reported variants for kinases were taken into account. The script generated a 20X20 matrix that gave a normalized count of each amino acid to other amino acids i.e., percent conversion of each amino acid. Normalized count = (Amino acid count in samples)/(Amino acid count from RefSeq)*100. This amino-acid exchange matrix was correlated with the chemical properties of mutating amino acids by analyzing the chemical shifts associated with variants among different populations and databases. The overall amino acid count for each class of amino acids was summed up for reference and alternate residues and the difference in the counts was called a chemical shift. The mutability of an amino acid is defined as the ratio of the total number of mutations for a specific amino acid in the data and the frequency of occurrence for that amino acid in the reference human genome. This mutational frequency was calculated for all the variants in IndiGen(AF >10%).

#### Statistical Analysis of amino acid Conversions

To determine statistically significant amino-acid exchanges among IndiGen and populations in 1000G data ( EAS, AMR, AFR, SAS, and EUR), one proportion z-test was used. Multiple hypothesis testing corrections (FDR) were performed using the Benjamini/Hochberg correction method. Corrected p-values of less than 0.05 were considered significant. (Supplemental Table S12).

#### Multiple Sequence Alignment and Protein Domain Analysis

To understand the effect of SNVs on protein’s function, it was checked whether the observed variation (SNVs) is conserved and falls under a protein domain or not. Clustal Omega [Sievers and Higgins, 2014] was implemented to perform the multiple-sequence alignment (MSA). The mutant protein sequence files in FASTA format were generated using a python script. For protein domain analysis, PfamScan [Madeira et al., 2019] web server maintained by EMBL-EBI was used. A single file of all protein sequences in FASTA format was provided as input (default parameters). It gave an output file consisting of a domain name, its start and end position corresponding to every input sequence (hmm name, hmm start, hmm end), and other information. Mutations observed within the domain region (hmm start - hmm end) were annotated as 0. For others, the distance of mutation from the domain region was also calculated.

#### Variant Protein Structure Generation

Computational protein structure prediction helps in generating a three-dimensional structure of proteins. The prediction is based on in-silico techniques and relies on principles from known protein structures primarily obtained by X-Ray crystallography, NMR Spectroscopy, and physical energy function. After applying the structure data filter, the native crystal structure corresponding to 12 genes in the IndiGen structure dataset were downloaded (PDB format) from RCSB Protein Data Bank. These native structures were considered as templates for mutant structure generation by mutating a single amino acid position from reference to alternative amino acid type for a particular gene/protein. This single reference amino acid of the protein was mutated using the rotkit function of PyMol [Schrödinger and DeLano] that allows access to its mutagenesis feature. Based on the requirements mentioned above, the protein’s crystal structure was downloaded from RCSB PDB and mutated using the rotkit function. This process was automated by python code. It was followed with energy minimization and refinement of these mutant structures (22 variants) using Chimera [Pettersen et al., 2004]. The parameters used for energy minimization include 1000 steepest descent steps with a step size of 0.02 Å and force-field AMBERff14SB. The impact of mutations on protein conformation, flexibility, and stability was predicted by Dynamut[Rodrigues et al., 2018]. The structural differences in native and mutant forms were analyzed using several tools like DSSP [Kabsch and Sander, 1983] for secondary structure annotation of mutated residue, HBPLUS [McDonald and Thornton, 1994] to study gain or loss of hydrogen bonds after the mutation and Naccess [Hubbard] to compare the solvent accessible surface area of the mutated residue.

#### Molecular Docking

Receptor-ligand docking was performed to analyze the effect of SNV in the binding affinity of the drug with its target protein before and after the occurrence of mutation. A set of kinase genes with FDA-approved drugs available in DGIdb were considered for this analysis. Only 10/12 genes (CHUK, EPHA7, GRK5, MAPK11, MAPK13, PI4K2B, PIK3CG, GRK4, TAOK3, and IRKA1) from IndiGen structure data were found to exhibit drug-gene interactions with 69 FDA-approved drugs (Supplemental Table S5). The protein structure files (in Protein Data Bank as a PDB format) for these ten genes and their 20 modeled variants were taken as receptors. Since our dataset comprised 69 ligands that were to be docked with 30 receptors, virtual screening was performed using AutoDock vina[Trott and Olson, 2009]. The structure files for ligands were downloaded from DrugBank [Wishart et al., 2008] and PubChem [Kim et al., 2019] in PDB format (Supplemental Table S6). Receptor preparation was performed by removing water molecules and heteroatoms, adding polar hydrogens, etc., followed by preparing ligands by assigning the correct AutoDock 4 atom types, adding Gasteiger charges, and detecting aromatic carbons. These prepared receptors and ligands were saved in PDBQT format. In the absence of any prior information about the target binding site, blind docking was carried out for all the protein-ligand pairs. The docking was performed to the center of the binding cavity using Cartesian coordinates that differed for every protein calculated using PyRx [Dallakyan, Sargis; Olson, 2015]. The docking grid with a dimension of 60 Å x 60 Å x 60 Å was used in each docking calculation with an exhaustiveness option of 100 (average accuracy). The maximum number of binding modes to generate was kept at 500 with an energy range of 20kcal/mol. Fifty iterations of these parameters for every target protein were followed.

#### Ligand similarity/diversity and Toxicity analysis

Bender et al. in their study established the correlation of chemical substructure with ADRs [Bender et al., 2007]. Yaminishi et al. studied about similar ADRs exhibited by chemically similar drugs [Yamanishi et al., 2010]. To identify chemically similar drugs with ADRs in our dataset, the ligand similarity and toxicity analysis was performed. The molecular similarity of the ligands (drugs) can be assessed using their structural features (e.g., shared substructures, ring systems, functional groups, topologies, etc.) of the compounds and their representations in the N-dimensional chemical space (Supplemental Table S7). These descriptors are often defined by mathematical functions of molecular structures. In this analysis, MACCS (Molecular ACCess System) keys with 166 keys and circular -Morgan fingerprints with radius 2 were used[Fernández-De Gortari et al., 2017]. These fingerprint-based similarity computations were implemented using the popular chemoinformatics package RDkit [Bento et al., 2020] in python. Tanimoto similarity coefficient was used to compute a quantitative score in order to measure the degree of ligand similarity and dissimilarity (1-similarity)-using weighted values of molecular descriptors. The toxicity of durgs were identified using ProTox II predictor [Banerjee et al., 2018].

#### Phenotypic drug-drug similarity

The tendency of a drug to bind to multiple targets is called drug polypharmacology; it is a well-known property of drugs. Reports have suggested the association of drug polypharmacology with the target protein family and binding site similarity of their primary targets [Jalencas and Mestres, 2013]. If two drug molecules target the same gene product, then they are expected to have similar activities and mechanisms of action[Prinz et al., 2016]. Thus, repurposed forms of similar drugs can act as alternatives to the ones with adverse drug reactions. Based on drug-gene interaction data obtained from DGIdb, several drugs were observed to have the same target protein (Supplemental Table S7).

## Results

### Indian variations in the kinome landscape

To first get an overview of the Indian variations present in the druggable kinome landscape, an exhaustive annotation of variation containing 545 kinase coding genes found in the IndiGen data and the families along with the number of drugs associated with them were mapped (Figure 1). It was observed that despite having more drug-gene interactions, very few genes from the atypical protein kinases family contained missense mutations. The SNVs in a conserved protein region can influence the protein structure and its stability and can affect the protein-protein or protein-drug binding affinity. A gene with more variation and multiple marketed drugs has a greater tendency of causing ADRs [Bachtiar and Lee, 2013]. It was found that the tyrosine kinase family, which has a maximum (1978) number of FDA-approved drugs, consists of the maximum (5013) number of variations. The class of kinases other than TK (Tyrosine Kinase) like the CMGC (cyclin-dependent kinase (CDK), mitogen-activated protein kinase (MAPK), glycogen synthase kinase (GSK3), CDC-like kinase (CLK), TLK (Serine/threonine-protein kinase tousled-like 1) and AGC (PKA, PKC, PKG) contain a large number of variations i.e., 10518, 1193, and 2943 respectively but the number of drugs with known Drug-Gene interactions were limited to 213, 185, and 339, which was comparatively less than the Tyrosine Kinase family. The CK1(casein kinase 1) class, among all others, contains the lowest (275) number of variations and lowest (18) drug-gene interactions. Kinase families associated with 545 kinase coding genes with the number of drugs and SNVs observed in each class are shown in Supplementary Table S11.

**Fig 1.**
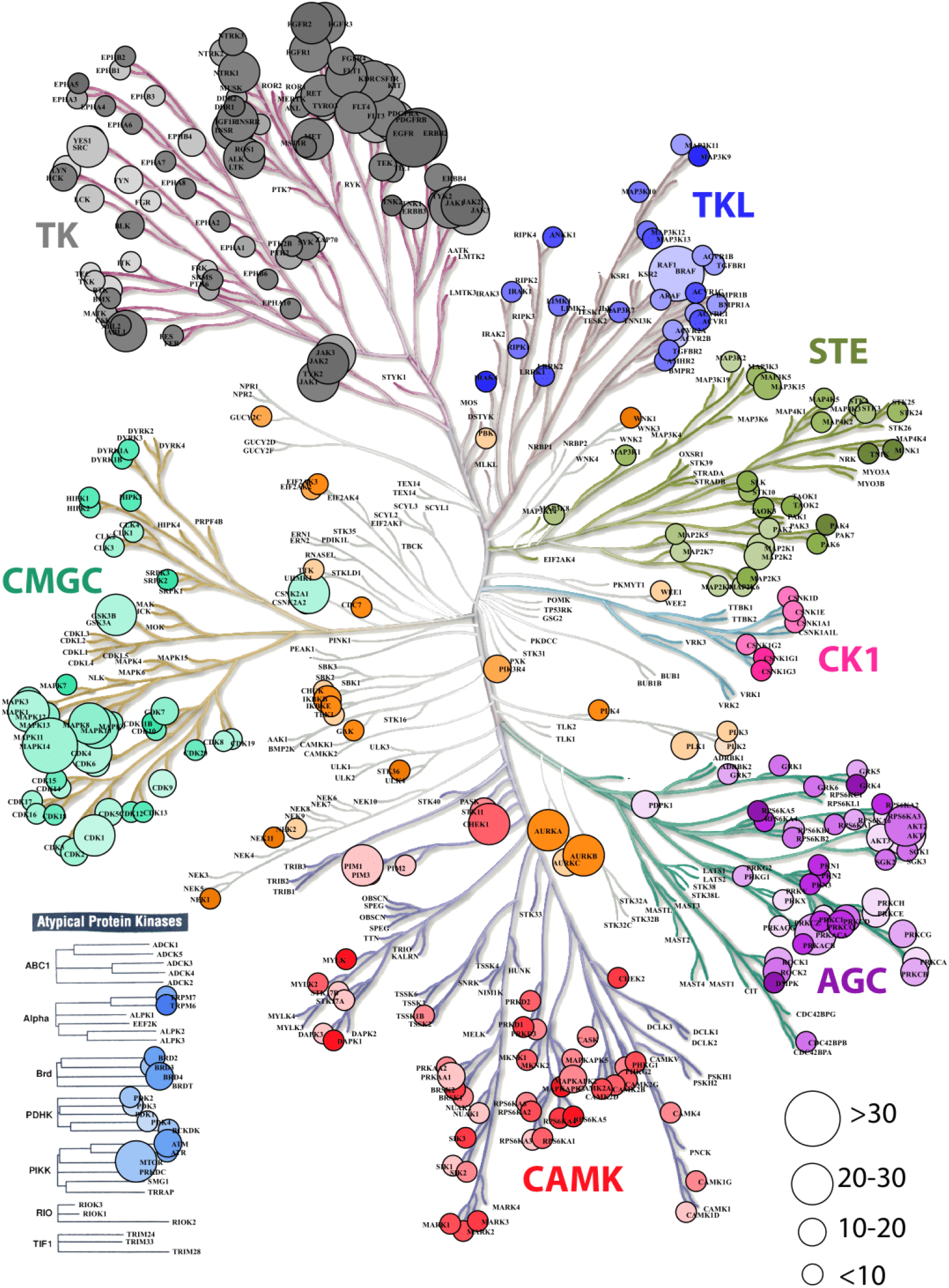
Dendrogram representation kinase coding genes in IndiGen data using KinMapbeta. The circle size represents the number of drug molecules available for a gene with known drug-gene interaction. The class of kinase is highlighted with a unique colour and the colour gradient in each data circle represents the number of variations present in IndiGen data for that gene.

### Analysis of amino acid changes of Druggable kinase genes among Indian and populations in 1000G data

The amino-acid mutation pattern in the Indian population was elucidated by generating an amino acid exchange matrix for all SNVs reported for 545 druggable kinase genes in IndiGen data. The number of times a specific amino acid has been converted to any other amino acid, resulting in the nsSNV was counted to understand how frequently a specific amino acid is mutating in a particular population. The AA-exchange frequency for every reference (as per RefSeq sequence) and alternative amino-acid pair (as per IndiGen data) i.e., was calculated and normalized in Figure 2A. Results from the analysis revealed that nearly 68% of Arginine(R) converts to Tryptophan (W), i.e., a hydrophobic amino acid converting to a basic polar amino acid. Similarly, 58% of Cystine (C) observed at reference SNV sites gets converted into Tyrosine (Y) i.e, a polar uncharged amino acid converting to a polar aromatic amino acid. Other amino acid conversions with moderate frequency (40-50%) were Leucine(L) to Phenylalanine(F), both non-polar amino-acids, Lysine(K) to Glutamic acid(E), which involved basic to acidic conversion, and Asparagine(N) to Aspartic acid(D), an amidic to acidic conversion. It was worth noticing that regardless of having a maximum number of codons (6) coding for Serine(S) and Leucine(L), the amino acid exchange for these two residues were comparatively lower than Tyrosine (Y) and Tryptophan (W), which have only one associated codon.

**Fig 2.**
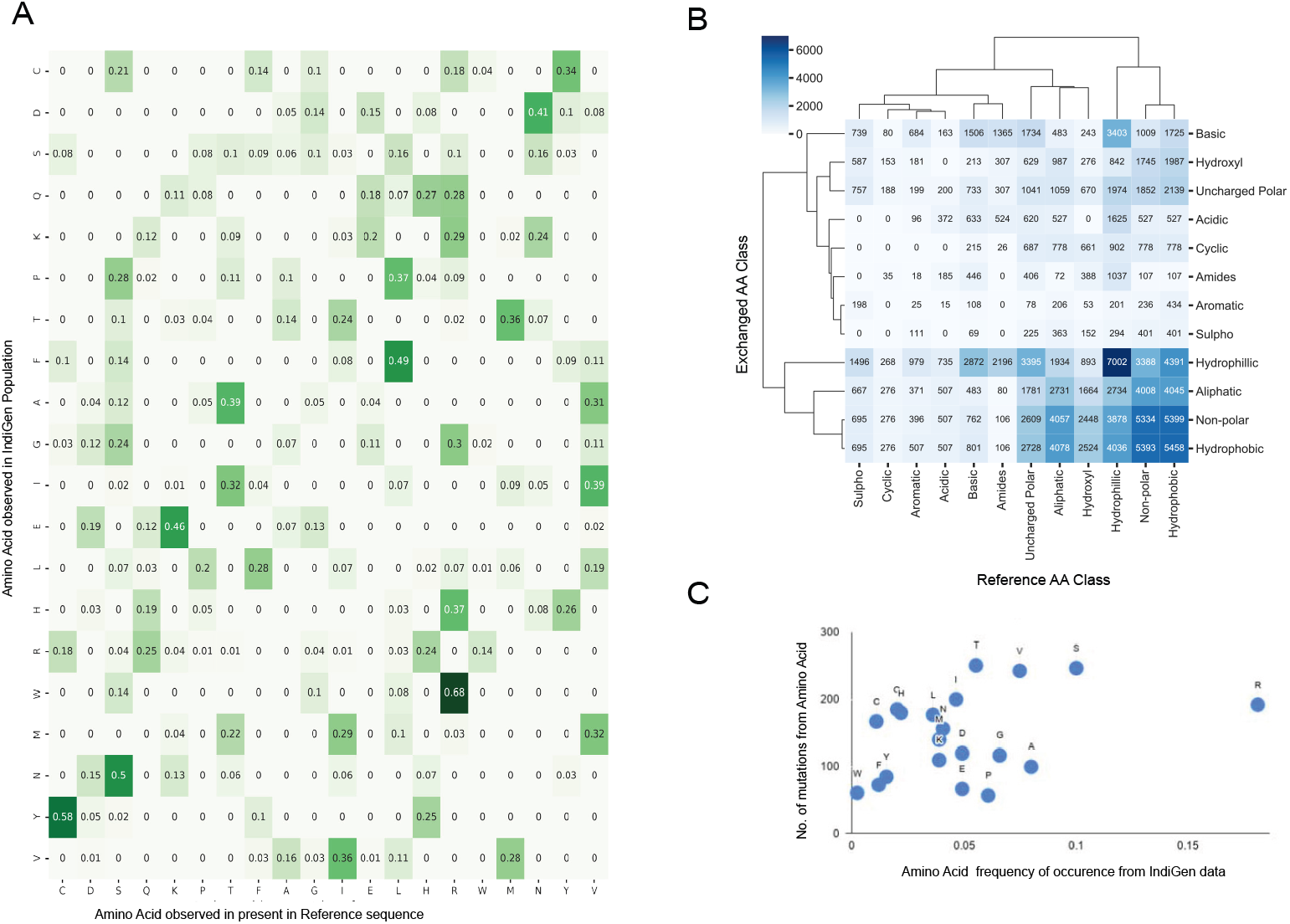
Sequence Analysis using amino-acid exchanges reported for 545 druggable kinase Coding genes in IndiGen Data: A. Amino-acid exchange matrix for reference and alternate amino acids of SNVs in IndiGen data. B. Clustermap showing chemical shift observed among the reference and alternate amino acids at SNV sites reported in IndiGen data. C. Scatter plot of mutability scores for each amino acid type in IndiGen data.

In order to comprehend the inter-conversion distribution of the chemical groups present in mutating amino acids and develop a coherent relation of these amino-acid conversions with physicochemical properties, a chemical shift analysis was performed. The mutating amino acids were classified on the basis of the nature of their side (R) groups into 12 chemical classes (Aliphatic, Hydroxyl, Cyclic, Aromatic, Basic, Acidic, Sulpho, Amides, Non-polar, Uncharged polar, Hydrophobic, and Hydrophilic). In Figure 2B, a cluster-map with 12 chemical classes on both axes is shown with values representing the number of amino-acids changing from one chemical class to another from reference (X-axis) to alternate sequence (Y-axis). Chemical groups with similar-level of changes in amino acids were clustered together. Intra-class conversions were observed for amino-acids belonging to hydrophilic (Ser, Thr, Tyr, Asn, Gln, Asp, Glu, Lys, Arg, His), hydrophobic (Gly, Ala, Val, Pro, Leu, Ile, Met, Trp, Cys, and Phe), and non-polar classes (Gly, Ala, Val, Pro, Leu, Ile, Met, Trp, Phe) supporting conservative replacement[French and Robson, 1983]. Many chemical groups have undergone more inter-class conversions than intra-class conversions, like Non-polar (Gly, Ala, Val, Pro, Leu, Ile, Met, Trp, Phe), Hydroxyl (Ser, Thr), Aliphatic (Gly, Ala, Val, Leu, Ile) and Uncharged polar groups (Ser, Thr, Cys, Tyr, Asn, Gln) have converted to Hydrophobic group. Similarly, amino acids from Amides (Asn, Gln), Basic (Lys, Arg, His), Acidic (Asp, Glu), Aromatic (Phe, Tyr, Trp), Cyclic (Pro), and Sulpho (Cys, Met) groups have mostly converted to Hydrophilic group.

In support of this, one more analysis was performed in which the reference amino acids were taken as per RefSeq hg38 sequence whereas altered amino acid at the same SNV site was taken from IndiGen data. These amino-acids were classified into six different chemical classes (Aliphatic (Gly, Ala, Val, Leu, Ile), Hydroxyl (Ser, Thr), Cyclic (Pro), Aromatic (Phe, Tyr, Trp), Basic (Lys, Arg, His) and Acidic (Asp, Glu)) to avoid any repetition of amino acids. The difference in amino-acid counts at the SNV site for each class was then plotted. A horizontal bar plot was generated with (Supplemental Figure S1A) six chemical classes of amino acids with respect to the amino acid counts in RefSeq(hg38) and IndiGen data. This chemical shift analysis confirms a net loss in basic, cyclic and aliphatic amino acid classes. In contrast, a net gain is observed in the hydroxyl, aromatic and acidic amino acid classes. It is important to note here that while the hydroxyl, Aromatic and Acidic amino acid class contains 2,3 and 2 amino acids, respectively, it contributes to the net gain; while the aliphatic class, with the maximum number of amino acids, showed a net loss in the amino acid count. This observation clarifies that the net gain or loss in any amino acid class is independent of its size.

In order to understand the relationship between the mutational frequency of specific amino acids with their frequency of occurrence in the IndiGen data, a mutability score for each amino acid type was calculated. In Figure 2C, mutability scores for amino acids observed in IndiGen data are shown. The plot shows that Arginine (R) is the most observed amino acid with >0.15 frequency of occurrence, whereas Tryptophan(W) is the least observed residue at the reference SNV site in IndiGen data. Amino acids like Valine, Serine, and Threonine have shown a greater propensity to get mutated than other amino acids. These observations are also in agreement with the inferences made from the amino acid exchange matrix(Figure 2A). In Figure 2A, Arginine(R) can be seen as the most mutable amino acid with the most significant amino-acid exchange frequency(maximum frequency - 0.68) and Tryptophan as the least mutable amino acid (maximum frequency-0.14).

After establishing an in-depth description of the Indian population, comparative sequence analysis was performed for the variants in IndiGen data with other populations, such as European (EUR), American (AMR), African (AFR), South Asian (SAS), and East Asian (EAS) populations from the 1000 genome data.

The count of each AA exchange from the RefSeq sequence to any other amino acid type in the alternate sequence for populations in 1000G data was calculated in a similar manner as was done for IndiGen data in Figure 2A. The difference in AA-exchange frequency pattern found in IndiGen with other populations in 1000G data was examined by performing a proportion z-test. The frequency of AA exchange for each reference to alternate AA pair (non-null, 144 Ref-Alt AA pairs) in IndiGen data was compared with its frequency in European (EUR), East-Asian (EAS), Ad Mixed American (AMR), African (AFR) and South-Asian (SAS) population in 1000G data (Supplemental Table S12). For every amino-acid exchange, a p-value and z-statistic were evaluated. The p-value obtained was adjusted, and a negative logarithm of this value was plotted in Figure 3A. In Figure 3A, a bubble plot is shown with reference AA and exchanged AA in the X and Y-axis respectively. The size of each bubble is inversely proportional to FDR corrected p-value, i.e., with the decrease in p-value the size of the bubble increases. The significant AA exchanges observed in IndiGen-EUR and IndiGen-SAS (for other population pairs- Supplemental Figure S1) population pair were highlighted in blue color. The number of statistically significant AA exchanges (FDR p-value < 0.05) between IndiGen and AMR, AFR, EUR, EAS, and SAS were 17, 16, 14, 12, and 5, respectively, suggesting that IndiGenic variations are more similar to the variations in the South Asian population as compared to others. The South-Asian population contains samples for Gujrati Indian from Houston (GIH), Punjabi from Lahor, Pakistan (PJL), Bengali from Bangladesh (BEB), Sri Lankan Tamil from the UK (STU) and Indian Telugu from the UK (ITU) due to which it shares a similarity with IndiGen data in terms of AA exchange frequency.

**Fig 3.**
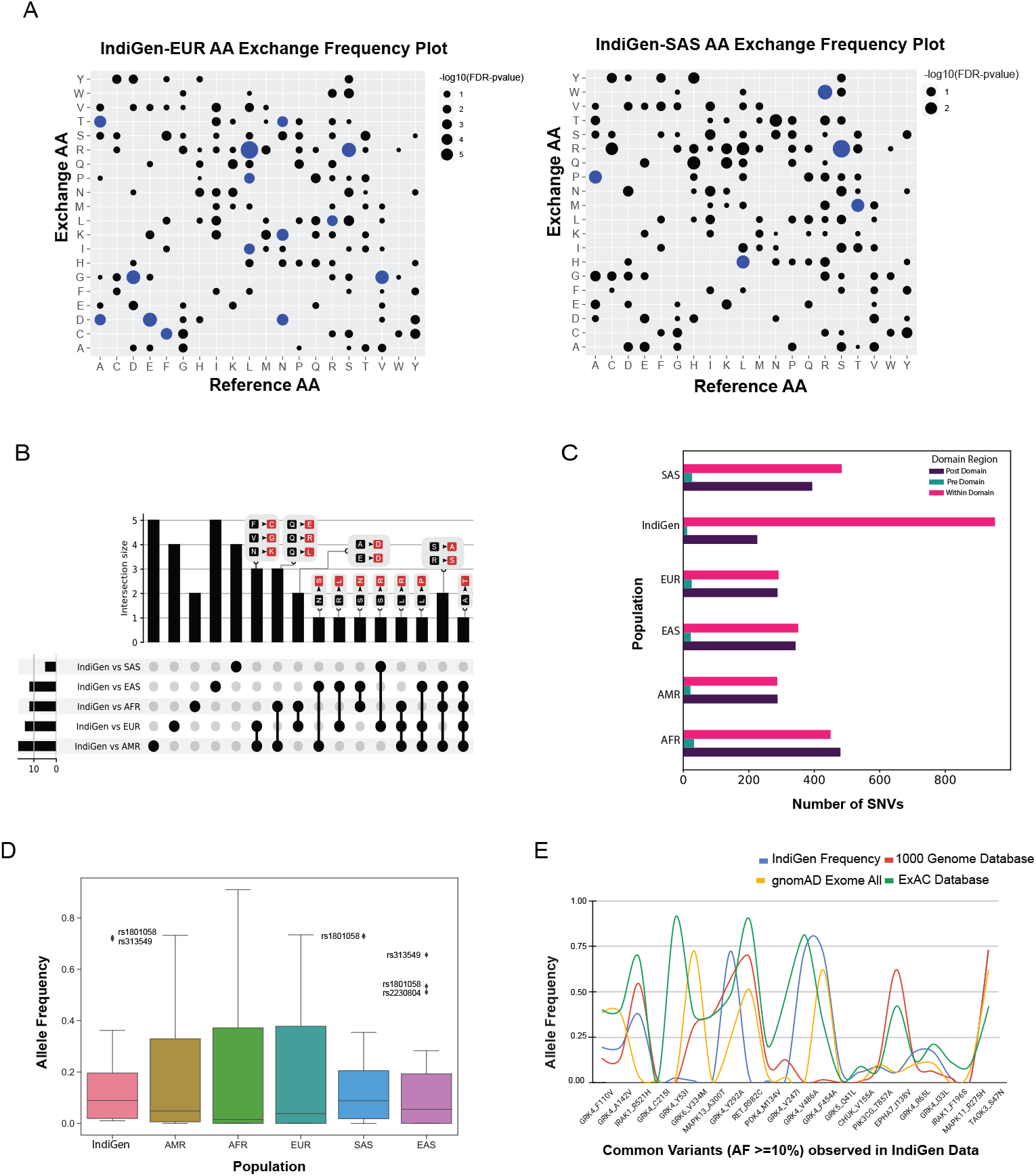
A. Comparing the trend of amino acid exchange among different populations from 1000 genome data with the Indian population. A. Bubble-plot was generated on the basis of the FDR corrected p-value associated with AA-exchange frequency for a particular Reference and Alternate AA observed in IndiGen data with EUR and SAS populations of 1000 genome data. The size of the bubble is proportional to the −log10 (p-value) linked with the amino acid exchange. AA exchanges with p-value < 0.05 are highlighted in blue color. B. An UpSet Plot of statistically significant AA exchanges observed in IndiGen w.r.t to other populations in 1000G data. C. A grouped bar plot showing the count of variations lying before (green), within (pink), and after the domain (violet) for variants in IndiGen and populations in 1000G data. D. A Box-plot for comparing allele-frequency distribution of common IndiGen variants (AF ≥ 10%) qualifying the filters used for structure data (22 variants) with different populations in 1000 genome data. E. IndiGen specific SNVs (22 variants in structure data) with AF ≥ 10% observed in different databases like 1000 genome project, gnomAD exome data, and ExAC database; with IndiGen variations on X-axis and their allele frequencies in different databases on Y-axis.

The amino-acid exchanges which were prevalent and specific across each population pair were studied by generating an UpSet plot (Figure 3B) using all the significant AA exchanges observed in Figure 3A. This UpSet Plot has four pieces, a bar plot (shows intersection size among the datasets), a graphical table below it (shows intersecting population pairs), common AA exchanges among the datasets shown above each bar) and a small bar chart left to the graphical table (shows the dataset size). The AA exchanges specific to each population, were represented by the bars above single dots in the graphical table. There were five AA conversions unique to IndiGen-AMR and IndiGen-EAS dataset; four conversions were unique in the IndiGen-SAS and IndiGen-EUR dataset, and two AA exchanges were unique to the IndiGen-AFR dataset. The AA exchange of Alanine to Threonine (A to T) was common in all the datasets except IndiGen-SAS. The conversion in amino acids common among three population pairs were Ser to Ala and Arg to Ser in IndiGen vs AFR, EAS and AMR, Leu to Pro in IndiGen vs EAS, AFR and AMR, and Leu to Arg in IndiGen vs AFR, EUR, and AMR (non-polar converted to the non-polar and basic group). Population pairs, IndiGen vs AMR and EUR and IndiGen vs AFR and AMR had a larger overlap among them with three common AA exchanges which were Phe to Cys; Val to Gly; Asn to Lys; and Gln to Glu; Gln to Arg; Gln to Leu (Uncharged polar converted to Acidic, basic and non-polar groups).

Upon having an in-depth understanding of the effects of variations on the sequence, we next explored the effect on the protein’s structure. Protein domain regions are stable conserved parts of a protein sequence and its 3D structure. Therefore, variants present in the protein domains are most likely to affect the protein structure, stability, and function. To determine the number of SNVs present in/out of a conserved protein domain, the protein domain analysis was executed. The variants in the IndiGen and 1000 genome data for European, American, African, East Asian, and South Asian populations were categorized into pre-domain, post domain, and within the domain regions, depending on the position of a variant. In (Figure 3C), a grouped bar chart was shown wherein X-axis six populations are there, and Y-axis represents the number of SNVs that fall inside (pink bars) or before/after a domain (green and violet bars). In all the populations, SNVs falling in pre-domain regions were less, suggesting that the populations from 1000 genome and IndiGen data revealed a larger bias for an SNV to fall within the protein domain or post-domain region as compared to the pre-domain region. The SNV count in the post and within domain regions were almost identical in EAS, AFR, AMR, and EUR variant data. In IndiGen, maximum variants (952) were falling within the domain while 226 variants were present in the post domain region, whereas only twelve variants were observed in the pre-domain region. Since, the occurrence of SNVs is more frequent inside the domain region, change in amino acid level can have a direct impact on protein’s structures, thereby on its function and stability too.

Reports have suggested the relationship between allele frequency and ethnicity of SNVs[Mattei et al., 2009, Mori et al., 2005]. The allele frequency distribution of common variants from IndiGen data (AF ≥ 10%, 22 variants in structure data) was compared with populations in 1000G data (Supplemental Table S3) in the Allele Frequency box plot shown in Figure 3D). The analysis revealed that the distribution of allele frequency didn’t vary much among the populations studied. The AF distribution of IndiGen, South Asian, and East-Asian populations were alike with very close median values and similar outliers rs1801058 and rs313549 belonging to GRK4 gene,i.e, Y292A, and V486A. A similar AF plot (Figure 3E) was generated using the same set variants wherein allele frequencies for common SNVs (22 variants in structure data) in IndiGen data were compared among other databases like 1000 Genome data, gnomAD exome database, and ExAC database (Supplemental Table S2).

### Structure level comparison of IndiGen and Disease-causing variants

To further understanding the SNV’s effect on the protein structures, IndiGen structure dataset was constructed by taking into account only variants of druggable kinases lying within the crystal length, thus giving only twelve kinase genes and corresponding 22 variants. Disease causing variants corresponding to these 12 genes were extracted from Humsavar data (217 variants) and compared (Supplemental Table S13). The structural characteristics like distribution of solvent-accessibility, secondary structure, conservation score and change in hydrophobicity of variants/variant residues in IndiGen structure data and Humsavar data were compiled and compared. For solvent accessibility comparison (in Figure 4A), a cutoff of 5% solvent exposure was applied onto the Naccess [Hubbard] results for variants in both datasets to distinguish between buried and exposed residues. It was observed that exposed residues are more prone to mutations in both the datasets. A similar observation was reported by Gong et al.[Gong and Blundell, 2010] in their work and they also stated that more than 60% of solvent exposed SNVs have a disease association. In IndiGen data, 81.8% of variant residues (22 residues) were found to be exposed which was roughly equal to solvent exposure of residues in Humsavar data with 81.1% exposed residues (74 residues). No significant difference was observed in solvent accessibility for variants in both datasets. The secondary structure preference of variants in both the datasets revealed that variant residues in IndiGen data have a slight preference to occur on alpha-helix part of the protein while the variants in Humsavar data share equal secondary structure preference for their occurrence either in alpha-helix or in loop/random coil of a protein (Figure 4B).

**Fig 4.**
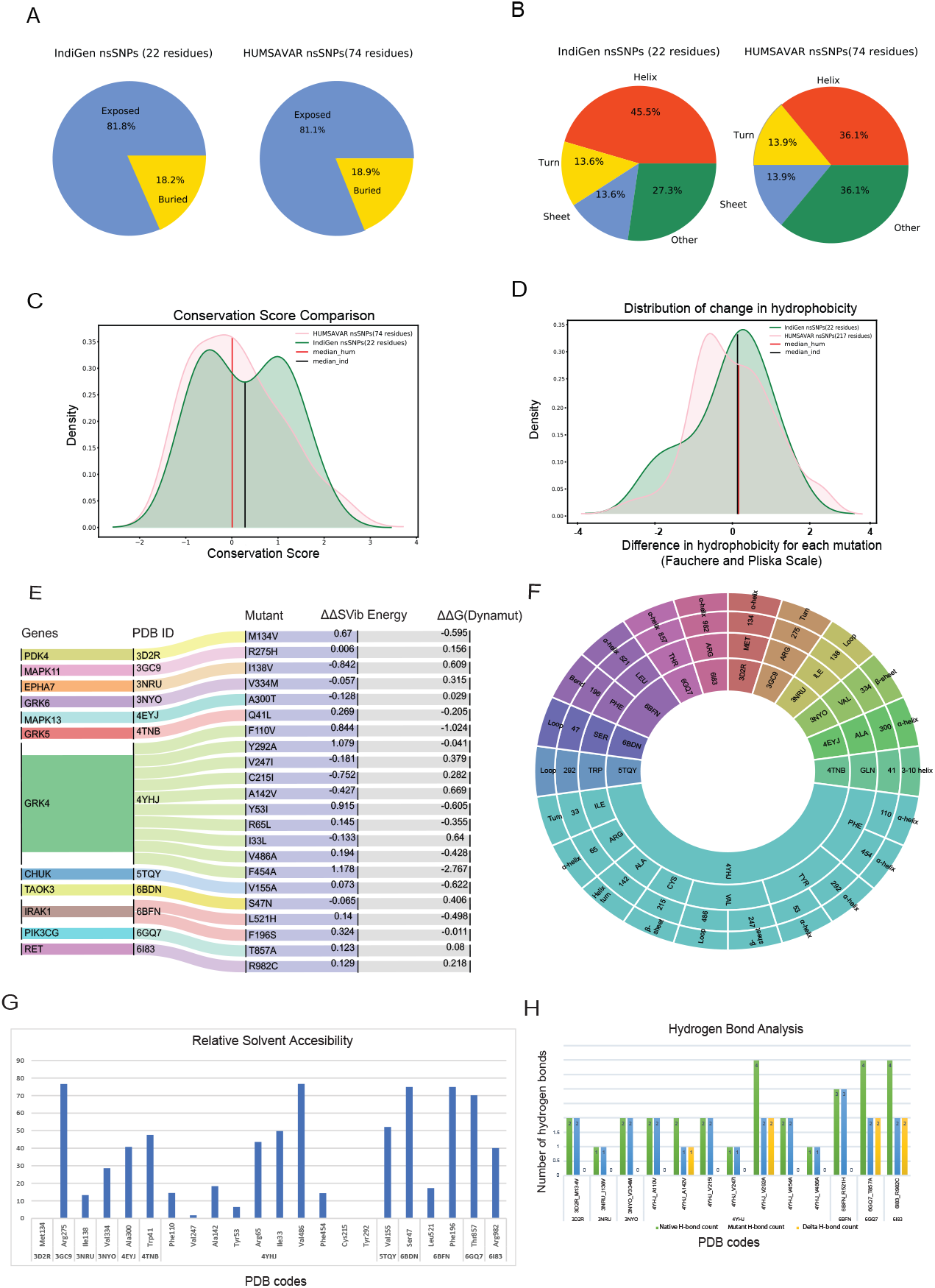
Comparison of structural characteristics of variants in IndiGen and Humsavar data: A. Solvent accessibility for the variants in both datasets. B. Secondary structure in which each of the variants occurs in both datasets. C. Conservation score and ∆Hydrophobicity distribution of variants in Humsavar and IndiGen data. D. The area under the curve present on the left of −2 (∆Hydrophobicity) belongs to the percentage of residues for which a significant increase in hydrophobicity after mutation was observed while decrease in hydrophobicity was observed for percentage of residue present on the right of +2 on X-axis. E. Alluvial plot representing change in folding energy (in kcal/mol) (*δ δ* G) and vibrational entropy by Dynamut energy for 22 variants. F. Sunburn Plot representing secondary structure assignment done by DSSP for mutant residues. G. Bar Plot showing relative solvent accessibility calculated by Naccess for mutated residues. H. HBPLUS results showing the number of hydrogen bonds made by mutated residue before mutation (green-bar), after mutation (blue-bar), and ∆H-bonds (yellow bars)

To study the evolutionary conservation of mutated residues, residue conservation scores for variants in IndiGen structure data (22 residues) and in Humsavar data (74 residues) were calculated using Consurf [Ashkenazy et al., 2016]. The distribution of conservation score for variants in both the datasets is shown in Figure 4C. The distribution followed by Humsavar dataset was nearly normal while the IndiGen curve follows a bimodal distribution with two peaks. Moreover, the median line divides the area under the curve into two equal halves. The median line for Humsavar data (0.007) was present closer to 0 than IndiGen data’s median (0.358). Hence, in order to elucidate the percentage of residues with more or less conservation, a threshold value of −1/+1 relative conservation score was considered. It was observed that the percentage of highly conserved residues (with Consurf conservation score greater than −1) was more in Humsavar distribution (steeper) than in IndiGen. Likewise, the percentage of highly variable residues (with conservation score >1) adhering to the area under the curve on the right of +1 was more for IndiGen data than for Humsavar data, indicating that Humsavar data has a higher percentage of residues that are involved in variations, being more conserved.

The distribution of change in hydrophobicity from reference to altered residue for variants in Humsavar and IndiGen structure data is shown in Figure4D. The medians for both the distributions were found next to each other and very close to 0, suggesting that the percentage of variations with increase or decrease in hydrophobicity is almost equal in both the datasets. In order to find out the percentage of residues with some significant change in hydrophobicity, a threshold value of −2 was considered for increase in hydrophobicity whereas +2 threshold was taken for decrease in hydrophobicity. It was observed that the percentage of varying residues with significant increase in hydrophobicity was observed for IndiGen structure data whereas the percentage of residues with significant decrease in hydrophobicity was found for Humsavar data.

### Effect of nsSNVs on structural properties of the protein

#### Structural stability of generated variants

Prior to investigation of the structural properties of nsSNVs in IndiGen structural data, impact of mutations on protein stability and flexibility was assessed using Dynamut [Rodrigues et al., 2018]. It performs normal mode analysis and follows a machine-learning algorithm to predict ∆ ∆G (change in folding energy, kcal/mol) and ∆ ∆ S (kcal/mol/K), Vibrational Entropy difference between native and mutant forms of a protein structure. The results from Dynamut revealed that 11/22 variants had ∆ ∆G negative suggesting destabilization after mutation and 14/22 variants had positive ∆ ∆ S indicating increase in structural flexibility after mutation, shown in Figure 4E. The plot shows 12 genes, their native protein structures codes (PDB IDs: 4YHJ, 5TQY, 3NYO, 6GQ7, 4TNB, 6BFN, 3GC9, 6BDN, 6I83, 4EYJ, 3NRU and 3D2R) and 22 mutants linked with their corresponding energy values. As per Dynamut predictions, a variants F454A and F110V of gene GRK4 (PDB ID: 4YHJ) has shown ∆ ∆G of −2.767 kcal/mol and −1.024 kcal/mol (Destabilizing) and change in vibrational entropy energy between wild-type and mutant (∆ ∆S-Vib) as 1.178 kcal/mol/K and 0.844 kcal/mol/K showing a high structural destabilization leading to increased molecular flexibility after mutation. The loss of aromaticity due to conversion of an aromatic amino-acid, Phenylalanine to aliphatic forms, Alanine and Valine in F454A and F110V variants could be reason behind instability of mutants, since aromatic rings are very stable and difficult to break thereby enhancing stability of the system (Supplemental Table S9).

#### Secondary Structure Annotation and Relative Solvent Accessibility of mutated residues

The secondary structure of a protein includes largely *α*-helix and *β*-pleated sheet structures, which is involved in local interactions between stretches of a polypeptide chain. The ability of a protein to interact with other molecules depends on amino acid residues located on the surface with high solvent accessibility. Any alterations in these residues may affect the protein’s functioning thereby increasing the importance behind the study of structural properties of mutated residues. Solvent accessibility (using Naccess [Hubbard]) and the secondary structure properties (using DSSP [Joosten et al., 2011]) of mutated residues were studied. The Figure 4F is a sunburn plot showing results for secondary structure assignment by DSSP. The plot consists of four concentric circles with innermost circle comprising 12 PDB IDs, second-inner circle comprising 3-letter code of reference amino acid present at mutant site, third-inner circle shows the mutant position and outermost circle contains the secondary annotation for that residue given by DSSP. The color coding was done on the basis of native PDBs. Majority of the variants were found to be present in alpha-helix region as compared to other regions of the protein.

In the Figure 4G, a bar plot is shown, wherein relative solvent accessibility (Y-axis) of mutated residues of 22 variants corresponding to 12 proteins (X-axis) are shown. The length of the bar represents the relative solvent accessibility scale associated with mutated residue and its position in the protein sequence. The relative solvent accessibility of two mutated residues belonging to PDB code 4YHJ (Y53I and C215I) was zero. The relative solvent accessibility of residues Arg275 and Val486 of mutants R275H (PDB ID: 3GC9) and V486A (PDB ID: 4YHJ) was greater than 75 suggesting that these two amino-acids are relatively more accessible than others. The results from this plot disclosed that there were 5 residues with more than 60 relative solvent accessibility (Arginine, Valine, Phenylalanine and Serine) belonging to 3GC9, 4YHJ, 6BDN and 6BFN PDB IDs (Supplemental Table S9).

#### Effect of nsSNV in hydrophobicity and hydrogen bonding

A single amino acid change may result in alteration of hydrophobicity or disruption of the hydrogen-bond network thus modifying the structure and function of the protein as well[Kumar and Biswas, 2019]. The change in hydrophobicity observed in mutants in IndiGen structure data were arranged according to Fauchere and Pliska scale [FAUCHÈRE et al., 1988] (Supplemental Figure S3A). In the IndiGen structure data, 12 out of the 22 variants exhibited decrease in hydrophobicity whereas an increase in net hydrophobicity was observed in the rest. The number of hydrogen bonds made by the alternate residue before and after the mutation were calculated using the HBPLUS program (Figure 4H). Variants 4YHJ A142V showed a loss of 1 hydrogen bond, while 4YHJ V292A, 6GQ7 T857A and 6I83 R982C resulted in loss of two hydrogen bonds.

#### Effect of nsSNV on ligand binding

Given the pharmacological importance of kinase proteins, molecular docking was performed to comprehend the effect of SNV in the drug-gene interaction. All FDA-approved drugs available in DGIdb for genes present in IndiGen structure data were docked against the native and mutant protein structures. The binding energy (also known as Gibbs free energy or ∆G in kcal/mol) between the receptor and ligand molecules is evaluated using AutoDock vina and compared among native and mutant docked complexes. It was found that in 45 out of 69 protein-drug pairs, change in binding energy ranges from 0.7 to −9.1 kcal/mol, whereas for the remaining pairs, no change in binding affinity was observed. Figure 5A represents the change in binding affinity observed for the 45 protein-drug pairs. In 32 protein-drug pairs, a decrease in binding energy was observed while 13 pairs have shown an increase in binding energy; indicative that the presence of an nsSNV destabilizes the complex. One protein-drug pair, T857A mutant of gene PIK3CG (PDB ID: 6GQ7), which when bound to drug Zinc sulfate (DrugBank id - DB09322) revealed a stark decrease in binding energy (−9.1 kcal/mol) when comparing the native-(−13.0 kcal/mol) versus mutant-(−3.9 kcal/mol) drug pair. Forty-five protein-drug pairs with differences in binding affinity were further considered for the binding site and ligand similarity (Supplemental Table S7).

**Fig 5.**
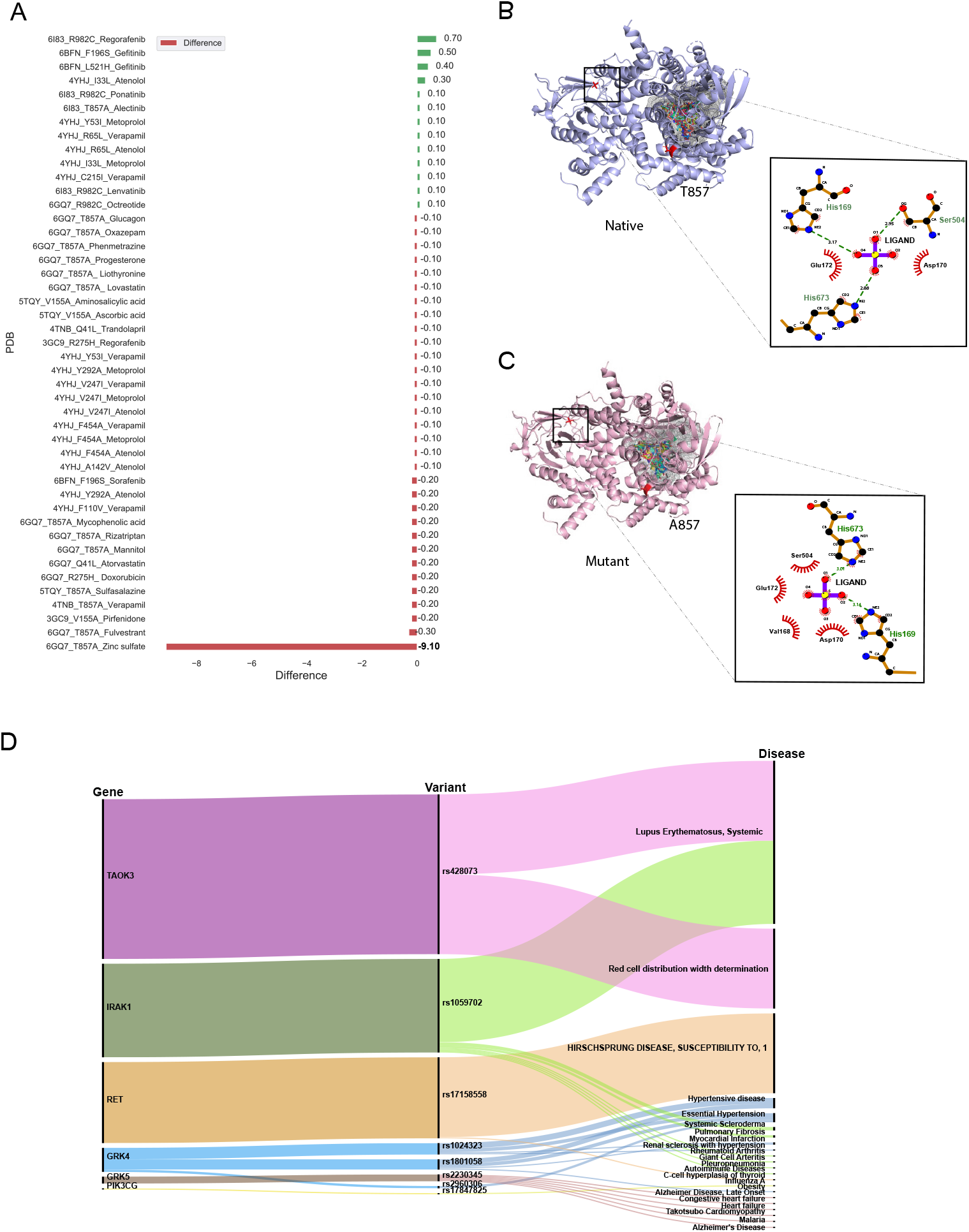
A. Bar plot showing docking results for 45 protein-drug pairs on x-axis and change in binding affinity observed on y-axis. Red bars represent a decrease in binding affinity and green bars represent increase in binding affinity after mutation. B. Ligand interaction diagram of native 6GQ7(PIK3CG gene) and its mutant T857A bound to Zinc Sulfate (DB09322) and main binding pocket (grey pocket) where majority of ligands docked.

It was observed that the binding pocket of the ligands in native and mutant forms for their respective receptors was the same, stipulating that presence of SNV didn’t change the binding site of drugs with their target protein. A snapshot of the first pose of ligand docked in the protein was taken in PyMol for all native protein-drug complexes (Supplemental Fig S2-(A-G)). The mutated residue in every complex is shown in red-color with sticks representation which was away from the binding pocket of the ligands in all cases(except in the case of 6GQ7-T857A). Ligand binding pockets (post docking) are shown in mesh representation with different colors in Supplemental Fig S2-(A-G).

In an attempt to find out the reason behind the huge decrease in binding affinity in the case of mutant T857A(PDB ID: 6GQ7)-zinc-sulfate(DrugBank ID - DB09322) complex the binding site residues of this drug in the native and mutant complex were compared and visualized in PyMol [Schrödinger and DeLano] and LigPlot+ [RA and MB, 2011], shown in Figure 5B,5C . It was observed that the location of the binding pocket-residues in mutant and native forms was unchanged, and the main binding pocket was away from the mutated residue. However, a decrease in one hydrogen bond was observed in the ligand interaction diagram of native and mutant complexes.

#### Gene, Variant and Disease association

Several databases like Online Mendelian inheritance in man (OMIM) [Amberger et al., 2011], disease-gene network (DisGeNET) database [Piñero et al., 2017], PharmGKB [Thorn et al., 2013] and Comparative Toxicogenomic Database (CTD) [Davis et al., 2019] could be used to retrieve diseases and/or ADRs associated with target genes or variants. DisGeNET is a database that collects information from Comparative Toxicogenomics DatabaseTM (CTDTM) [Davis et al., 2019], UniProt[Consortium, 2021], Orphanet [Rath et al., 2012], the Mouse Genome Database (MGD) [Eppig et al., 2015] and the Rat Genome Database (RGD) [Shimoyama et al., 2015] for finding gene-disease relationship and ClinVar [Landrum et al., 2016], the NHGRI-EBI GWAS Catalog [Welter et al., 2014], and the GWAS db [Becker et al., 2004] for variant-disease association. It assesses the relation between gene/variant with a disease using Gene-Disease-Association (GDA) score or Variant-Disease-Association score (VDA). The score ranges from 0 to 1 on a scale of 1 to 10 and depends on the number and kind of sources/publications (degree of curation, organisms) supporting the association. The list of rsIDs of variants in the structure dataset was given as input to the DisGeNET webserver, and the summary of results obtained is shown in Figure 5D. It is an alluvial plot with genes, variants, and disease-associated, where the thickness of the variant-disease line is proportional to the VDA score obtained from DisGeNET. Variants rs428073, rs1059702, and rs17158558 were reported to be linked with a risk to Systemic Lupus Erythematosus, risk of association with hematological traits, and HIRSCHSPRUNG disease with a VDA score equivalent to 0.7, indicating at least one curated source in support of this variant-disease association. The effect of genetic variation in the drug-response present in our structure dataset was investigated using the PharmGKB [Thorn et al., 2013] tool. The gene-set in our structure data were found to be interacting with 69 drugs as per DGIdb and there were 22 variants associated with it. The phenotypic information due to the association of these genes, drugs, and variants was collected from PharmGKB data. It was observed that pharmacogenetic variants, rs1024323 and rs1801058 ( F110V, A142V, Y292A, V486A, and F454A) evoked a phenotypic effect (Hypertension, Nephrosclerosis and Kidney issues) on the interaction between GRK4 gene and Metoprolol drug (Supplemental Table S10).

#### Ligand Similarity/diversity and Toxicity Analysis

The 45-protein drug pairs with delta binding energy observed after docking were considered for this analysis. In total there were seven different PDB structures (6GQ7, 5TQY, 3GC9,4TNB, 6I83) with sixteen respective mutations and 28 drugs as shown in (Supplemental Table S7). All drug-like chemicals from our ligand dataset were considered for chemical similarity analysis. From this analysis, it was observed that all the associated drugs exhibit a great molecular diversity (Figure 6). The maximum pairwise similarity for Morgan2 fingerprints and MACCS fingerprints has a Tanimoto score of 0.40 and 0.70, respectively. On the other hand, the pairwise dissimilarity (1-similarity) for Morgan2 fingerprints and MACCS fingerprints has a Tanimoto score of 0.98 and 0.90, respectively. The computational prediction platform ProTox-II [Banerjee et al., 2018], which includes cheminformatics-based machine learning models for predicting 46 toxicity endpoints, was used to predict toxicity profiles of compounds/drugs. For the prediction of various toxicity endpoints, such as acute toxicity (LD50 values), hepatotoxicity, cytotoxicity, carcinogenicity, mutagenicity, immunotoxicity, adverse outcomes pathways (Tox21), and toxicity targets, ProTox-II [Banerjee et al., 2018] integrates many statistical methodologies such as molecular similarity, pharmacophores, and fragment propensities, as well as machine learning models (off-targets). In vitro assays (e.g. Tox21 assays, Ames bacterial mutation assays, hepG2 cytotoxicity assays, Immunotoxicity assays) and in vivo cases were used to create the predictive models (e.g. carcinogenicity, hepatotoxicity).

**Fig 6.**
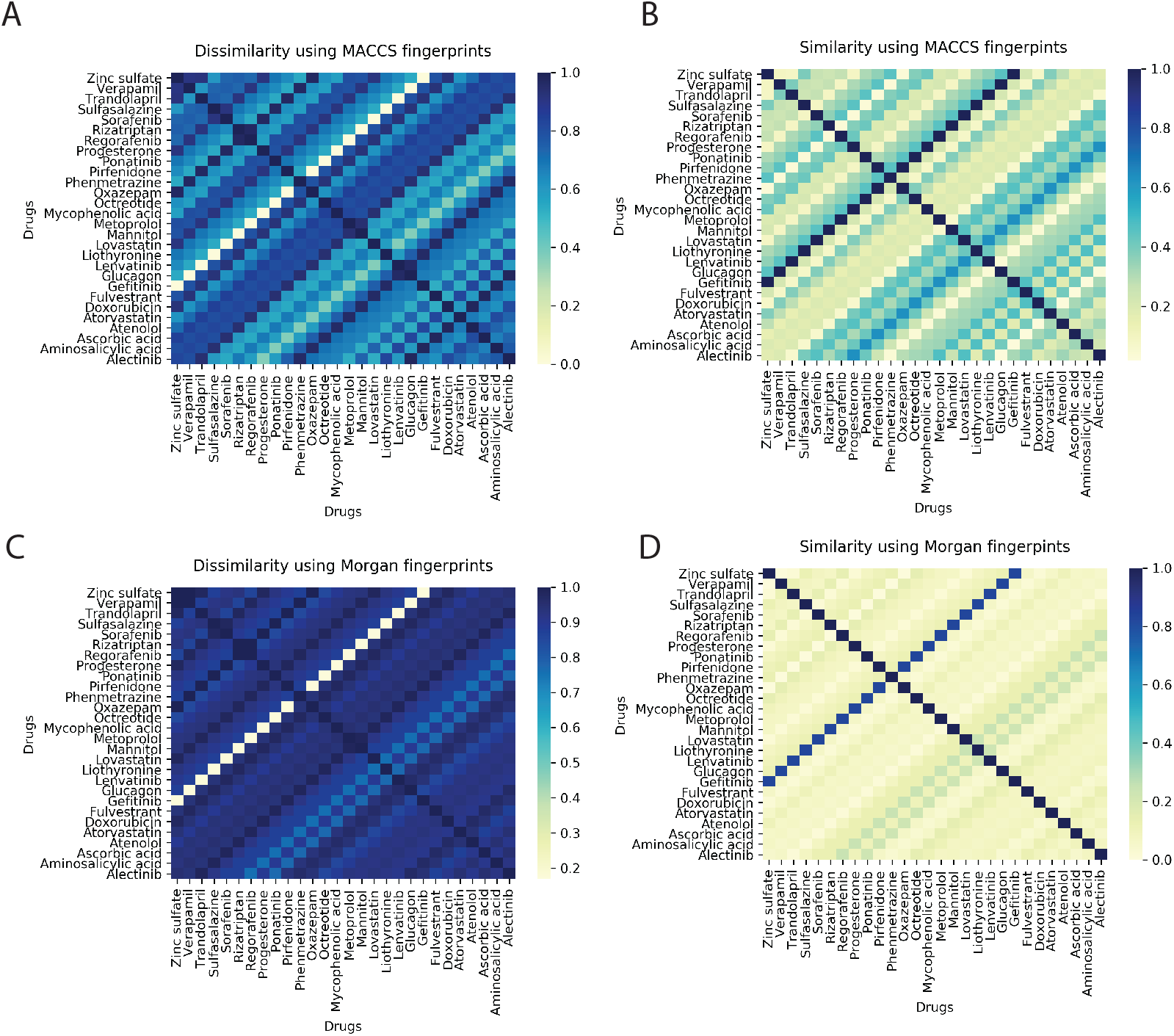
A. Heatmap showing pairwise ligand dissimilarity using MACCS fingerprints. B. Heatmap showing ligand similarity using MACCS fingerprints C. Ligand dissimilarity using Morgan fingerprints. D. Heatmap showing ligand similarity using Morgan fingerprints. Similarity and dissimilarity (1-similarity) score is represented using Tanimoto coefficient (taking a value between 0 and 1, with 1 corresponding to maximum similarity)

As per the predictions made by ProToxII (Supplemental Table S8), it was observed that the mycophenolic acid (DB01024) is an immunosuppressant drug which was predcited to be hepatotoxic, immunotoxic, and cytotoxic and was found to have an interaction with gene PIK3CG (PDB ID:6GQ7,Mutant T857A). It also inhibits SR-MMP(mitochondrial membrane potential) with a confidence score of 0.79. Another interesting observation was the drug Regorafenib (DB08896) which was also predicted to be hepatotoxic and was active in two different stress response pathways SR-MMP and SR-p53. Regorafenib is associated with adverse events like hypertension, stomatitis, the abnormal liver function [Krishnamoorthy et al., 2015]. However, the exact mechanism of developing hypertension is not very well-defined. Abnormalities in liver function were also reported in the case of Regorafenib[De Wit et al., 2014]. The drug progesterone (DB00396) was predicted to be active in six adverse outcome pathways(AOPs). Like progesterone, many other drugs can result in such molecular inhibition/ activation of NR-AR by progesterone, and can result in reduced AR signaling/impaired follicle recruitment as cellular or tissue level response and may be impaired fertility in organism[Pivonello et al., 2020]

#### Phenotypic drug-drug similarity

A single drug can bind to multiple proteins, similarly, a single protein (molecular target) can interact with several drugs. These kinds of interactions between molecular targets and drugs can result in therapeutic effects along with clinically relevant adverse effects [Kanji et al., 2015]. The analysis of the drug-target interactions is an important step in the discovery of additional applications of the drugs already approved in the market, also called- drug repurposing, and in drug safety through the explanation of undesirable adverse effects caused by drug administration.

In order to look for phenotypically similar drugs in IndiGen data a list of protein IDs and drug molecules associated with them was considered (Supplemental Table S7). This information could be useful to get insights into similar drugs present in IndiGen structure data. A correlogram was plotted with drug names on the x/y axis. The positive and negative correlation was shown by blue and red color circles. The color intensity depends on the correlation coefficient. (Supplemental Fig S3-B). A strong correlation (more blue dots) can be observed from this plot indicating the promiscuous nature of drugs (binding to multiple targets) or target proteins. For instance drugs, Fulvestrant and Rizatriptan are chemically dissimilar (similarity score 0.20 in Figure 6). However, in terms of phenotypic drug-drug similarity - they are highly similar as they bind to the same protein target highlighting the differential binding ability of kinases to a set of fairly specific inhibitors. Another interesting example drug pair that is evident from the Figure 6 and Supplemental Figure S3-B) are the drugs Metoprolol (*β*1 receptor blocker) and Atenolol (beta-blocker), where both these drugs have a ligand similarity of Tanimoto Coefficient of 0.20. However, the same drugs are similar in terms of their phenotypic profiles and have a correlation coefficient of 1. Interestingly, both the drugs have similar ADRs profile- renal and urinary disorders and hypertension [Charles and Ferris, 2020, Manuscript, 2013] . This is also documented by the RWE data from EU ADRs reporting system (Supplemental Table S14). Both the drugs are associated with the gene GRK4 (Supplemental Table S7) and several studies have been published in literature supporting genetic variation in GRK4 gene and drug-induced hypertension [Frey et al., 2017] and renal disorders [Armando et al., 2015, Sanada et al., 2016] . Understanding the role of genetic variants related to blood pressure regulations and salt sensitivity may reveal new therapeutic drug targets and optimise the therapeutic effects of the drugs in the Indian population. Furthermore, Metoprolol is known to be a poor metabolizer of CYP2D6 which results in higher systemic concentrations [Banerjee et al., 2020]. Several studies have been reported on the importance of CYP2D6 polymorphisms on the therapeutic responses of cardiovascular presents treated with beta-blockers [Rau et al., 2002].

## Discussion

Adverse drug reactions are often associated with genes that are more prone to variations and targeted by multiple drugs [Wilke et al., 2007]. To have an understanding of distribution of variations in IndiGen data in kinoe landscape, the kinome dendrogram for all the druggable kinase genes was constructed (Figure 1). This revealed that the tyrosine kinase class consisted of a large number of variations and was found to be associated with numerous drugs. Receptors tyrosine kinases (RTKs) are involved in a broad range of functions such as proliferation, differentiation, and apoptosis of cells and have been extensively used as drug targets in cancer studies. Many of the tyrosine kinase inhibitors are antibody-based drugs used in the treatment of tumors, malignancies, and inflammatory diseases[Bennasroune et al., 2004]. In chemical shift analysis the intra-class conversions of hydrophilic and non-polar AA classwas observed. The conservative mutations of such kind can affect the protein’s stability which can modulate its functioning and catalytic pattern followed by it in different organisms[Rodriguez-Larrea et al., 2010]. Studies have shown there is a strong correlation between the frequency of occurrence of amino acids in the human genome and the number of associated codons[Alwi, 2005]. On the contrary, observations made in the amino acid exchange matrix and chemical shift analysis(Figure 2) suggested that mutation from one amino acid type to another was independent of the number of codons coding for any amino acid. It was also found that amino-acids have greater tendency to convert into hydroxyl, aromatic and acidic amino acid classes. The mutability plot (Figure 2D) revealed that Arginine (R) is more mutable than other amino acids and the probable reason behind this could be the presence of CpG dinucleotide in the codons coding for Arginine which is relatively vulnerable to mutations[de Beer et al., 2013].

Ancestry has a very important role to play in the evolution of an SNV in different ethnic groups of a population. This also indicates that there is a relationship between allele frequency and ethnicity of the population. Even a fractional exchange of amino acids can have a completely different impact on different populations. The comparative study of amino acid exchange frequency of IndiGen with other populations in 1000 Genome data stipulated (Figure 3B) samples in AMR data have a significant difference with IndiGen variations. In the contrary, variations in SAS population share a significant similarity with IndiGen data. An amino-acid exchange Alanine to Threonine (A to T) was statistically significant and found to be common in all the datasets except IndiGen-SAS suggesting the similarity of Indian and South-Asian genetic variations.

Some variants were found to be common (high AF) in the Indian population and rare (low AF) in other populations (population-specific variants) indicating that it will be affecting the Indian population with higher frequency than others(Figure 3D). On comparing allele frequency of Indian mutations with the ones present in publicly available databases it was inferred that many conserved mutations in IndiGen data are still understudied as none of the existing databases contains these mutations (referring to IndiGen data=samples from 1000 individuals of strict Indian ethnicity)(Figure 3E). Protein domain regions are stable conserved parts of a protein sequence and its 3D structure. Therefore, variants present inside the protein domains are more likely to affect the protein structure, stability and function. The comparative study of variants on the basis of their position with respect to domain location suggested that many Indian variants were present either within the domain or in the post-domain region.(Figure 3C)

One of the most useful predictors of the phenotypic effects of missense mutations is protein structural information and stability. Missense mutations can disrupt protein structure and function in one of two ways: they can destabilise the entire protein fold or they can change functional residues, such as active sites or protein-protein interactions, and pathogenic mutations are enriched in both the buried cores of proteins and in protein interfaces[Gerasimavicius et al., 2020]. Reports have claimed that buried amino acids are often observed to be associated with diseases and commonly observed in functional sites [Iqbal et al., 2020]. On the contrary, in relative structural analysis of IndiGen and Humsavar dataset, it was found that residues with relatively higher solvent accessible surfaces were more prone to mutations (Figure 4A)

Mutations that occur in a properly structured part of a protein are more likely to be pathogenic than mutations that do not, due to their strong destabilizing effect on protein structure. According to stability analysis performed by Dynamut, 11 variants were found to destabilize protein’s structure and from 11 destabilizing variants, 7 were found to be present in the helix region of the protein. IndiGen variants occur more in the alpha-helix region while Humsavar variants share equal secondary structure preference for their occurrence either in alpha-helix or in loop/random coil of a protein (Figure 4B). Several studies have suggested that secondary structure elements like sheets and helices vary a lot in their ability to tolerate mutations. This differential tolerance of mutations could be due to a difference in a number of non-covalent residue interactions within these secondary structure units.[Abrusán and Marsh, 2016]. The conservation score distribution implied a higher percentage of residues with greater conservation in that Humsavar data than in IndiGen data. Since Humsavar variants are reported to be associated with a disease it is highly likely that their presence in highly conserved regions could be a reason behind their disease occurrence. Hydrophobic interactions and hydrogen bonds are the two most prevalent interactions present in protein structure. Hydrophobes as the name suggests tend to isolate themselves from water molecules due to which many hydrophobic amino acids are often found to be buried inside the protein structure. Contrasting results were observed in hydrophobicity distribution with the significant increase in hydrophobicity for varatioans in IndiGen structure data whereas a decrease in hydrophobicity was found for Humsavar variant data. (Figure 4C)

Occurence of nsSNVs at the ligand-binding sites (LBSs) can influence protein’s structure, stability and binding affinity with small molecules. Interesting findings claimed that ligand binding residues have a significantly higher mutation rate than other parts of the protein [Kim et al., 2017]. In order to validate whether a single amino acid substitution can change the binding affinity of a ligand with its target protein or not, molecular docking of ligands(FDA-approved drugs) with native and mutant structure was performed. The docking results suggested that since the mutated residue was away from the binding pocket not much difference in binding affinity was observed in native and mutant forms except in T857A mutant in which a polar amino acid has converted to a non-polar amino acid leading to loss of two hydrogen bonds (4H), thereby decreasing the binding affinity of ligand(Zinc-sulphate) with protein. Moreover, the molecular diversity of 12 drugs binding to 6GQ7 (PIK3CG) suggest the promiscuous nature of the kinase and enable insights which are relevant for understanding polypharmacology and negative side-effects. Further analysis of these and other inhibitors that bind to PIK3CG, clustered by phenotype information, can give us deeper insights into targeted kinase inhibitor design. Additionally, structural dissimilar drugs can share similar ADRs because of their strong phenotypic drug-drug similarity as explained in the case of Metoprolol and Atenolol. Both these drugs interact with the genetic variants of the GRK4 gene. Understanding the role of genetic variants related to blood pressure regulations and salt sensitivity may reveal new therapeutic drug targets and optimize the therapeutic effects of the drugs in the Indian population. Within the scope of this study, we have investigated and analyzed several factors that can contribute to ADRs. From chemical space, to molecular target space including genetic variants and their relation to specific ADRs. ADRs can be related to on-target interactions, or related to the complexity of genome regulation and the heterogeneity of the particular ADRs. As a perspective, our approach can be helpful in identifying new ADRs and understanding the mechanism behind existing ADRs and can guide further experimental studies.

While in this study, we have explored common variants present in the Indian population, sampling lower allele frequencies shall be also useful, in the future, to understand the underlying fundamentals of rare diseases. Additionally, experimental validation of the findings in this study shall provide further credence to the results. This study on IndiGen variant data may assist in redesigning the healthcare system from “One Size Fits for All” to “Population or Individual Specific Drug System” and a big step towards the effective treatment of patients due to utilisation of drugs with fewer side-effects. Furthermore, a database of ADRs reporting systems needs to be established to understand the risk associated with multiple therapies resulting in drug-drug interactions and to safeguard the health of the Indian population. The analysis presented in this study, supporting the screening and detection of the ADRs specific to the Indian population will certainly be more meaningful when it is directly compared with known ADRs data obtained for the Indian population. Currently, this preliminary study may be helpful in devising strategies for the pre-clinical and post-market screening of drug-related ADRs in the Indian population. The European Medical Agency (EMA) and FDA (The United States Food and Drug Administration) has a current reporting system for ADRs, and this real-world evidence data (RWE) are important to contribute to the knowledge of where, when, and how ADRs take place. The lack of awareness of ADRs and missing reporting system on Indian ADRs which is publically accessible is a major obstacle to the completeness of studies specific to Indian ADRs. Integrating the RWE data along with the poly-pharmacological and poly-toxicological data would pave the way for effective medicine prioritizing patient safety.

## Author contribution

G.P., N.M., A.R. conceptualized the study, performed analysis and wrote the manuscript. D.S., R.C.B., A.J., M.I., V.S., M.K.D., A.M., S.S., and V.S. generated the IndiGen data and assisted in the inputs in the manuscripts. P.G. performed the domain analysis. G.P., N.M., and P.B. performed the pharmacogenomic analysis and P.B., S.S. and V.S. gave critical insights during the manuscript writing.

