## Supplementary Material for "Comprehensive assessment of Indian variations in the druggable kinome landscape highlights distinct insights at the sequence, structure and pharmacogenomic stratum"

### SUPPLEMENTARY FIGURES

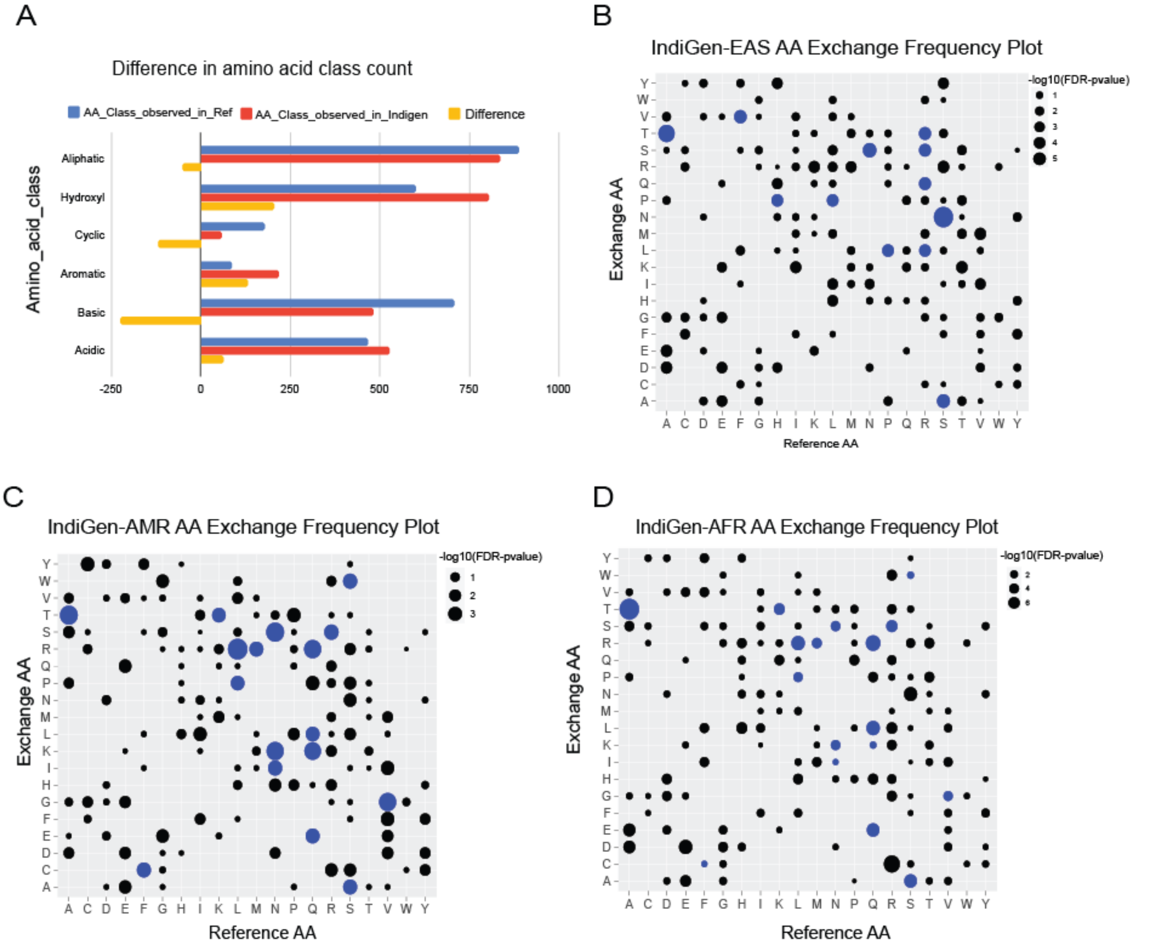


**Supplemental_Figure_S1:** ***A***. *Chemical changes observed from reference amino acid (RefSeq(hg38)) to alternative amino-acids at SNP sites reported in IndiGen data.* ***B,C,D***. *Bubble-plot was generated*

*on the basis of the FDR corrected p-value associated with AA-exchange frequency for a particular Reference and Alternative AA observed in IndiGen data with EAS , AMR and AFR populations of 1000 genome data.*

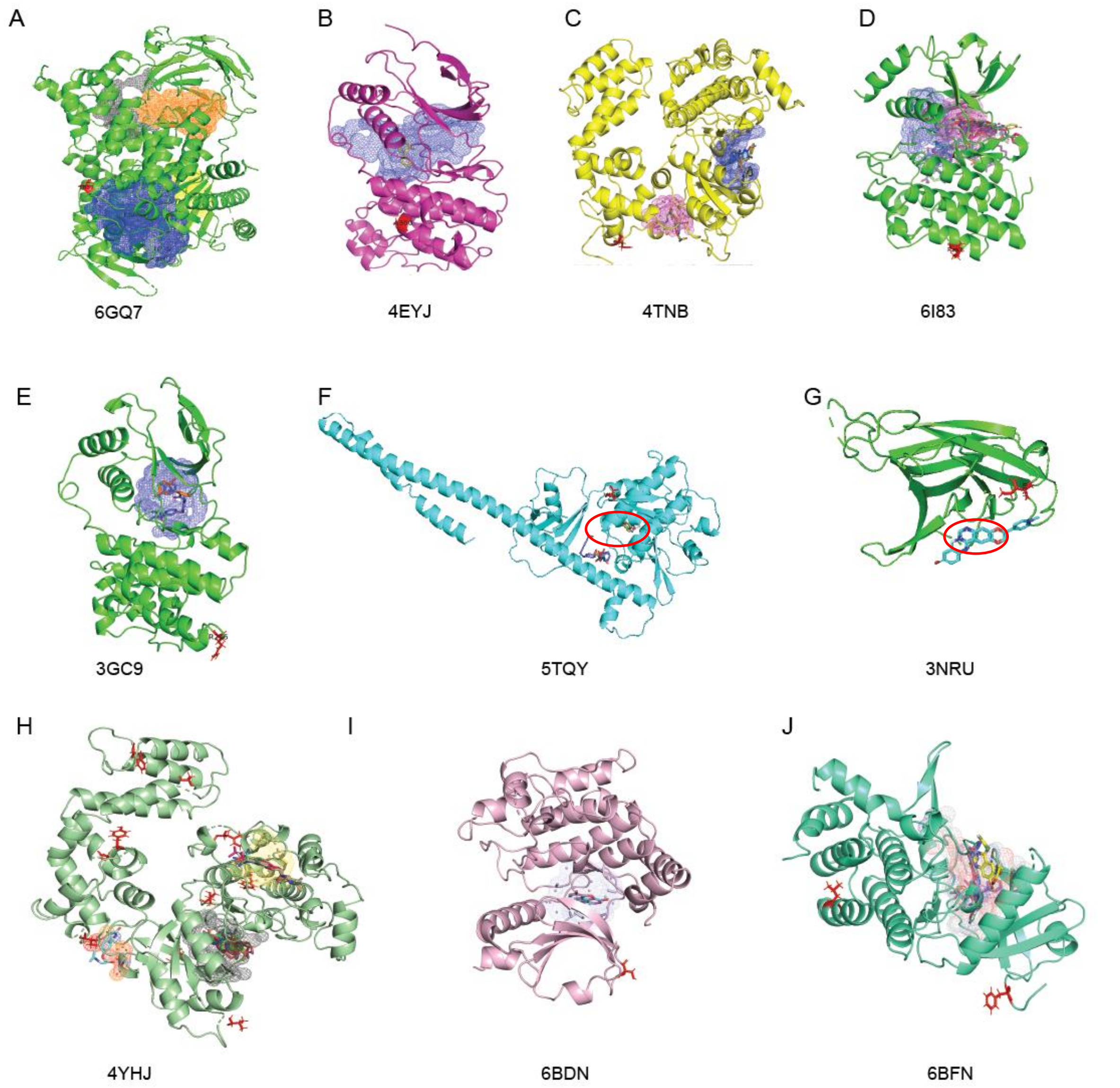


**Supplemental_Figure_S2**. (**A-G**). *Snapshot of docked complexes A. 6GQ7 docked with 34 ligands bound at 4 different pockets (grey, blue, orange and yellow color), mutated residue Thr at 857th position shown in red stick representation. B. 4EYJ docked to 1 ligand (blue pocket). C. 4TNB docked with 4 ligands in two pockets (pink, blue color). D.6I83 docked with 15 ligand molecules in two pockets (purple and pink). E. 3GC9 docked with 2 ligands in one pocket(blue). (F) 5TQY docked with 5 ligands in one pocket (red circle). (G) 3NRU docked with 1 ligand in one pocket (red circle) (H) 4YHJ docked with 4 ligands in 3pockets (grey, orange, yellow) (I) 6BDN docked with 1 ligand a pocket (purple) (J) 6BFN docked with 2 ligands in 1 pocket (orange)*

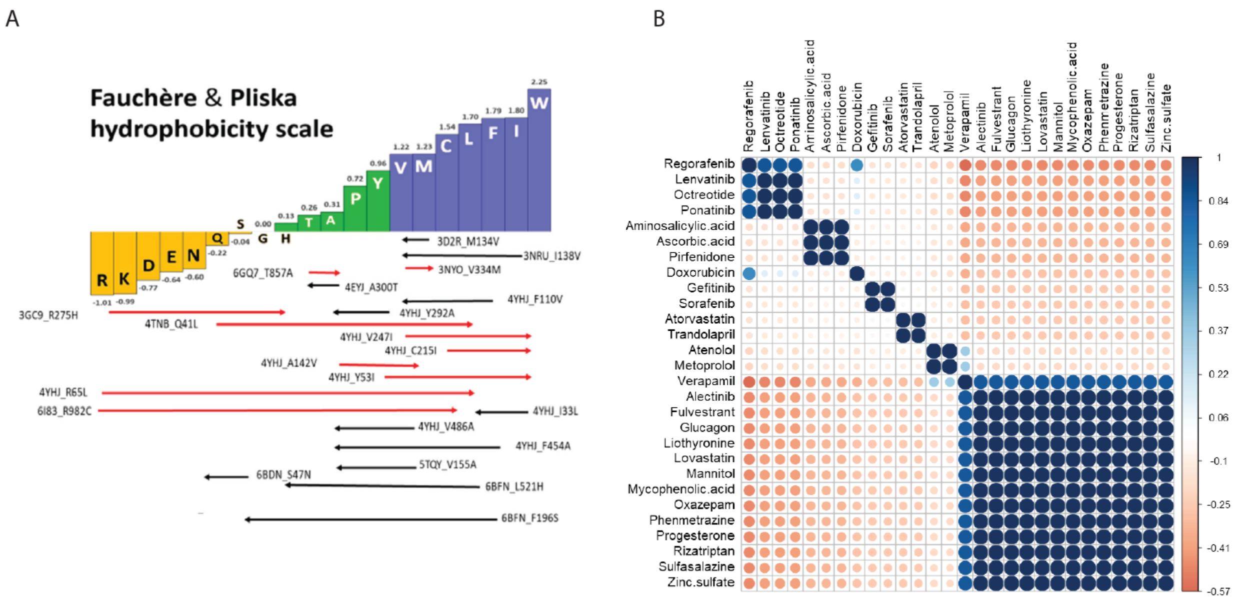


**Supplemental_Figure_S3**. ***A***. *Fauchere and Pliska hydrophobicity scale showing change in hydrophobicity observed in 22 mutations (red-line for increase in hydrophobicity and black line for decrease in hydrophobicity).* ***B.*** *Phenotypic drug-drug correlogram*

### SUPPLEMENTAL_TABLES

**Supplemental_Table_S1: 545 druggable kinase coding genes for the study.**

| Gene name | | | | | |
|---|---|---|---|---|---|
| TGFBR2 | TBK1 | ABL2 | RET | IRAK3 | MAP3K14 |
| INSRR | IRAK4 | BRD2 | CHEK1 | ERN1 | STRADA |
| BMPR1A | MAP3K13 | CAMK2B | AURKA | ERN2 | STRADB |
| ACVR1B | MAPKAPK3 | CDK8 | AKT1 | MAP4K1 | TAF1L |
| EPHA1 | MAPKAPK5 | CDK19 | MTOR | MAP4K2 | TAOK1 |
| EPHA4 | STK3 | GCK | JAK2 | MAP4K3 | TTBK1 |
| EPHA5 | STK4 | CHUK | INSR | LRRK1 | TTBK2 |
| EPHA6 | NEK2 | CHEK2 | ABL1 | LIMK2 | TBCK |
| EPHA10 | PAK2 | FRK | FGFR2 | MARK4 | TESK1 |
| EPHB3 | PAK4 | PTK2B | PDGFRA | MAST1 | TESK2 |
| EPHB6 | PDK2 | GSK3A | PDGFRB | MAST2 | TSSK2 |
| TIE1 | PDK4 | PRKCI | KIT | MAST3 | TSSK3 |

| | | | | | |
|---|---|---|---|---|---|
| PRKAA2 | PLK2 | MAPK9 | PIK3CD | MAST4 | TSSK4 |
| PRKAB2 | PLK3 | ARAF | FGFR1 | MASTL | TSSK6 |
| CSRP2 | PRKD2 | MAP2K6 | FLT4 | MELK | MLKL |
| CSRP2P1 | PRKG1 | TGFBR1 | FLT1 | CAMKK1 | TLK1 |
| BRD1 | RPS6KA6 | ROS1 | PIK3CB | CAMKK2 | TLK2 |
| BRSK1 | PTK6 | ATR | SRC | MKNK2 | TRIB1 |
| CAMK1G | SRPK2 | PRKCZ | FLT3 | MAP3K20 | TRIB2 |
| CAMK2D | MAP3K1 | MAPK13 | MET | MOS | TRIB3 |
| CASK | MAP3K5 | PRKD1 | MAPK14 | MAP4K4 | KALRN |
| CLK1 | MAP3K8 | MAP2K3 | BRAF | NRK | OBSCN |
| CLK3 | TAF1 | MAP2K4 | PIK3CG | MYLK3 | SPEG |
| MATK | BMX | MAP2K5 | KDR | MYLK4 | TRIO |
| STK17A | TGFBR3 | MAP2K7 | ERBB2 | TTN | TWF1 |
| STK17B | ACVR1 | PRKAA1 | CDK2 | AAK1 | TWF2 |
| DYRK1A | CAMK4 | BCR | EGFR | STK16 | ULK1 |
| FES | CSNK1A1 | PRKCH | PIK3CA | LATS1 | ULK2 |
| DMPK | CSNK1G2 | PTK2 | GUCY2D | LATS2 | ULK3 |
| CDC42BPB | CSNK1G3 | MAPKAPK2 | BMPR2 | STK38 | ULK4 |
| GRK1 | CSNK1E | BLK | BMPR1B | NIM1K | VRK1 |
| GRK4 | CDK3 | EPHB4 | MUSK | NEK3 | VRK2 |
| GRK6 | CSK | AURKC | ROR1 | NEK4 | VRK3 |
| GRK7 | DAPK1 | BTK | ROR2 | NEK5 | PKMYT1 |
| HIPK1 | HASPIN | NTRK3 | PTK7 | NEK6 | WEE2 |
| HIPK2 | IKBKE | PRKCG | LTK | NEK7 | WNK2 |
| HIPK3 | ITPKA | PRKCQ | AATK | NEK8 | WNK3 |
| ITPKB | MYLK2 | PDPK1 | LMTK2 | NEK9 | WNK4 |
| IRAK1 | NUAK1 | HCK | LMTK3 | NEK10 | STK32A |

| | | | | | |
|---|---|---|---|---|---|
| MAP4K5 | EIF2AK2 | CDK7 | RYK | MYO3A | STK32B |
| LRRK2 | RPS6KB2 | CDK9 | STYK1 | MYO3B | STK32C |
| LIMK1 | PDK3 | MAPK3 | PIP5K1A | SBK1 | STK25 |
| MAP3K12 | PLK4 | PRKCE | PIP5K1C | SBK2 | STK26 |
| MARK1 | PRKAR1A | TYK2 | PIP4K2B | SBK3 | TTK |
| MARK2 | PRKACB | ROCK2 | PIP4K2C | PINK1 | FGFR3 |
| MARK3 | PKN2 | YES1 | ADCK2 | PDIK1L | IRAK2 |
| MKNK1 | SIK2 | TEK | ADCK1 | STK35 | MAP3K19 |
| MAP3K10 | SPHK2 | PIK3C2A | COQ8A | TEX14 | RPS6KB1 |
| MAP3K11 | MAP3K7 | PIK3C2B | FOXN3 | NLK | PRKD3 |
| RPS6KA5 | TEC | PIK3C2G | ADCK5 | NRBP1 | ANKK1 |
| RPS6KA4 | TXK | FYN | TNK1 | NRBP2 | RPS6KA1 |
| MINK1 | ACVRL1 | PRKCD | PRKAG1 | NUAK2 | FGR |
| TNIK | ACVR1C | MAPK11 | PRKAG2 | BMP2K | GSK3B |
| MYLK | MERTK | ROCK1 | PRKAG3 | ALPK1 | PIK3R1 |
| NEK1 | TNK2 | LYN | BRD7 | ALPK3 | IGF1R |
| NEK11 | CAMK2G | NTRK2 | BRD8 | CIT | NTRK1 |
| GAK | CSNK1D | CDK6 | BRD9 | EIF2AK1 | MAP2K1 |
| PAK6 | CDK20 | MAPK12 | BUB1 | STKLD1 | CSF1R |
| PAK5 | CDK13 | PIK3C3 | BUB1B | DSTYK | CDC42BPA |
| EIF2AK3 | GRK5 | PIK3R4 | TP53RK | STK31 | TNNI3K |
| PI4K2A | MAP3K9 | PIK3R6 | CAMK1 | UHMK1 | HIPK4 |
| PI4K2B | PIM2 | BRD4 | PNCK | PAK1 | HUNK |
| PHKG1 | PIM3 | IKBKB | CAMK2A | PAK3 | ILK |
| PHKG2 | CDK11B | PLK1 | CAMKV | PASK | ITPKC |
| PRKACG | PKN1 | AKT3 | STK33 | PIP5K1B | SRPK1 |
| PRKX | SIK1 | PRKCB | STK40 | PIP4K2A | SRPK3 |

| | | | | | |
|---|---|---|---|---|---|
| PKN3 | SIK3 | CDK5 | CSNK1A1L | CDK11A | MAP3K3 |
| BCKDK | SGK1 | STK11 | CSNK2B | PRKAR1B | MAP3K4 |
| RIPK1 | SLK | PRKCA | CERK | PRKAR2A | MAP3K6 |
| RPS6KA2 | SPHK1 | SYK | CLK2 | PRKG2 | MAP3K15 |
| SGK2 | CDK14 | FGFR4 | CLK4 | PRPF4B | PRKDC |
| STK10 | CDK16 | PIM1 | CDKL1 | PSKH1 | MAPK7 |
| MAP3K2 | CDK17 | RAF1 | CDKL2 | PSKH2 | CAMK1D |
| TAOK2 | CDK18 | AKT2 | CDKL3 | KSR1 | CSNK1G1 |
| TAOK3 | GUCY2C | MAP2K2 | CDKL4 | KSR2 | CSNK2A2 |
| TSSK1B | EPHA2 | ERBB4 | CDKL5 | ICK | DAPK3 |
| PBK | MST1R | CSNK2A1 | DCLK1 | MAK | DYRK1B |
| STK36 | BRD3 | MAPK8 | DCLK2 | MOK | PI4KB |
| WNK1 | CDC7 | ERBB3 | DCLK3 | RIPK2 | BRSK2 |
| STK24 | CDK10 | ATM | DAPK2 | RIPK3 | ITK |
| ACVR2A | TRPM6 | LCK | DYRK4 | RIPK4 | WEE1 |
| ACVR2B | TRPM7 | MAPK1 | DYRK2 | RIOK1 | PIK3R3 |
| AMHR2 | CDK12 | JAK1 | DYRK3 | RIOK2 | AURKB |
| EPHA3 | PRKAR2B | JAK3 | EEF2K | RIOK3 | EIF2AK4 |
| EPHA7 | RPS6KA3 | PRKACA | MAPK15 | RPS6KC1 | CDC42BPG |
| EPHA8 | ZAP70 | MAPK10 | MAPK4 | RPS6KL1 | SMG1 |
| EPHB1 | CDK15 | ALK | MAPK6 | SCYL1 | SNRK |
| EPHB2 | AXL | CDK1 | FER | SCYL2 | PI4KA |
| TYRO3 | DDR2 | CDK4 | OXSR1 | PKDCC | SRMS |
| DDR1 | PRKAB1 | PIK3R5 | STK39 | SGK3 | PIK3R2 |
| PXK | STK19 | | | | |

**Supplemental_Table_S2: Allele Frequency of variations observed in IndiGen data vs other databases.**

| Gene | Variation | Indigen frequency | 1000 genome database | Genome AD Exome All | ExAC database | avsnp |
|---|---|---|---|---|---|---|
| GRK4 | F110V | 0.196 | 0.136 | 0.3833 | 0.4016 | rs1024323 |
| GRK4 | A142V | 0.196 | 0.136 | 0.3833 | 0.4016 | rs1024323 |
| IRAK1 | R521H | 0.378 | 0.546 | 0.036 | 0.693 | rs1059702 |
| GRK4 | C215I | 0.027 | 0.021 | 0.011 | 0.0068 | rs1140085 |
| GRK4 | Y53I | 0.027 | 0.001 | 0.027 | 0.91 | rs1140085 |
| GRK6 | V334M | 0.012 | 0.313 | 0.724 | 0.373 | rs143935970 |
| MAPK13 | A300T | 0.011 | 0.373 | 0.009 | | rs144262262 |
| GRK4 | Y292A | 0.724 | . | . | 0.487 | rs150897108 |
| RET | R982C | 0.036 | 0.693 | 0.514 | 0.901 | rs17158558 |
| PDK4 | M134V | 0.011 | 0.106 | 0.043 | 0.201 | rs17847825 |
| GRK4 | V247I | 0.027 | 0.126 | 0.03 | . | rs1801058 |
| GRK4 | V486A | 0.724 | 0.003 | 0.002 | 0.812 | rs1801058 |
| GRK4 | F454A | 0.724 | 0.017 | 0.621 | 0.329 | rs2230345 |
| GRK5 | Q41L | 0.074 | 0.002 | 0.003 | 0.014 | rs2230349 |
| CHUK | V155A | 0.06 | 0.009 | 0.002 | 0.089 | rs2278107 |
| PIK3CG | T857A | 0.089 | 0.101 | 0.099 | 0.07 | rs28763991 |
| EPHA7 | I138V | 0.058 | 0.622 | 0.058 | 0.421 | rs2960306 |
| GRK4 | R65L | 0.171 | 0.116 | 0.101 | 0.121 | rs2960306 |
| GRK4 | I33L | 0.171 | 0.063 | 0.106 | 0.212 | rs2960306 |
| IRAK1 | F196S | 0.026 | 0.026 | 0.001 | 0.11 | rs33932986 |
| MAPK11 | R275H | 0.01 | 0.01 | 0.072 | 0.101 | rs41270090 |
| TAOK3 | S47N | 0.731 | 0.731 | 0.621 | 0.421 | rs428073 |

**Supplemental_Table_S3: Allele frequency Indian v/s other populations from 1000 genome data(1000g2015).**

| Gene | avsnp150 | AFR | EUR | SAS | EAS | AMR | Indigen |
|---|---|---|---|---|---|---|---|
| CHUK | rs2230803 | 0.001 | 0 | 0.048 | 0.055 | 0.001 | 0.06 |
| EPHA7 | rs2960306 | 0.537 | 0.378 | 0.155 | 0.094 | 0.329 | 0.171 |
| GRK4 | rs1024323 | 0.634 | 0.402 | 0.174 | 0.19 | 0.383 | 0.196 |
| GRK4 | rs1801058 | 0.91 | 0.57 | 0.728 | 0.534 | 0.643 | 0.724 |
| GRK4 | rs2960306 | 0.537 | 0.378 | 0.155 | 0.094 | 0.329 | 0.171 |
| GRK5 | rs2230349 | 0.002 | 0.076 | 0.205 | 0.283 | 0.095 | 0.074 |
| GRK6 | rs143935970 | 0 | 0 | 0.012 | 0 | 0.006 | 0.012 |
| IRAK1 | rs1059702 | 0.967 | 0.853 | 0.393 | 0.22 | 0.574 | 0.411 |
| IRAK1 | rs33932986 | 0.015 | 0.022 | 0.018 | 0 | 0.033 | 0.01 |
| MAPK11 | rs41270090 | 0 | 0 | 0.006 | 0 | 0.003 | 0.175 |
| MAPK13 | rs144262262 | 0 | 0.011 | 0.026 | 0 | 0.01 | 0.011 |
| PDK4 | rs17847825 | 0.007 | 0.11 | 0.239 | 0.193 | 0.048 | 0.216 |
| PIK3CG | rs28763991 | 0.077 | 0.038 | 0.088 | 0.032 | 0.084 | 0.089 |
| RET | rs17158558 | 0.011 | 0.016 | 0.033 | 0.032 | 0.023 | 0.036 |
| TAOK3 | rs428073 | 0.745 | 0.683 | 0.759 | 0.714 | 0.761 | 0.731 |

**Supplemental_Table_S4: IndiGen Structure Data- consisting of 12 genes and their 22 variants**

| Native | Chain | Variation |
|---|---|---|
| 3D2R | A | M134V |
| 3GC9 | A | R275H |
| 3NRU | A | I138V |
| 3NYO | A | V334M |
| 4EYJ | A | A300T |
| 4TNB | A | Q41L |
| 4YHJ | A | F110V |
| 4YHJ | A | Y292A |
| 4YHJ | A | V247I |
| 4YHJ | A | C215I |
| 4YHJ | A | A142V |
| 4YHJ | A | Y53I |
| 4YHJ | A | R65L |
| 4YHJ | A | I33L |
| 4YHJ | A | V486A |
| 4YHJ | A | F454A |
| 5TQY | A | V155A |
| 6BDN | A | S47N |
| 6BFN | A | L521H |
| 6BFN | A | F196S |
| 6GQ7 | A | T857A |
| 6I83 | A | R982C |

**Supplemental_Table_S5: Table with gene names, PDB ids and observed mutations in IndiGen data and no. of FDA-approved drugs given by DGIdb for these genes.**

| Gene | PDB | Variant | No. of drugs |
|---|---|---|---|
| MAPK11 | 3GC9 | R275H | 2 |
| EPHA7 | 3NRU | I138V | 1 |
| MAPK13 | 4EYJ | A300T | 1 |
| GRK5 | 4TNB | Q41L | 4 |
| CHUK | 5TQY | V155A | 5 |
| RET | 6I83 | R982C | 15 |
| PIK3CG | 6GQ7 | T857A | 34 |
| GRK4 | 4YHJ | F110V | 3 |
| | | Y292A | |
| | | V247I | |
| | | C215I | |
| | | A142V | |
| | | Y53I | |
| | | R65L | |
| | | I33L | |
| | | V486A | |
| | | F454A | |
| TAOK3 | 6BDN | S47N | 1 |
| IRAK1 | 6BFN | L521H | 3 |
| | | F196S | |
| 10 genes | 10 PDBs | 20 variants | 69 drugs |

**Supplemental_Table_S6: Data used for docking**

| Native | Variation | DrugBank Id |
|---|---|---|
| 3GC9 | R275H | DB04951 |
| 3GC9 | R275H | DB08896 |
| 3NRU | I138V | DB05294 |
| 4EYJ | A300T | DB04951 |
| 4TNB | Q41L | DB00519 |
| 4TNB | Q41L | DB00661 |
| 4TNB | Q41L | DB00999 |
| 4TNB | Q41L | DB00335 |
| 5TQY | V155A | DB00795 |
| 5TQY | V155A | DB00244 |
| 5TQY | V155A | DB00126 |
| 5TQY | V155A | DB00233 |
| 5TQY | V155A | DB06151 |
| 6GQ7 | T857A | DB00091 |

| | | |
|---|---|---|
| 6GQ7 | T857A | DB00104 |
| 6GQ7 | T857A | DB00227 |
| 6GQ7 | T857A | DB00279 |
| 6GQ7 | T857A | DB00363 |
| 6GQ7 | T857A | DB00388 |
| 6GQ7 | T857A | DB00396 |
| 6GQ7 | T857A | DB00481 |
| 6GQ7 | T857A | DB00641 |
| 6GQ7 | T857A | DB00655 |
| 6GQ7 | T857A | DB00742 |
| 6GQ7 | T857A | DB00830 |
| 6GQ7 | T857A | DB00842 |
| 6GQ7 | T857A | DB00947 |
| 6GQ7 | T857A | DB00953 |
| 6GQ7 | T857A | DB00984 |
| 6GQ7 | T857A | DB00997 |
| 6GQ7 | T857A | DB01024 |
| 6GQ7 | T857A | DB01064 |
| 6GQ7 | T857A | DB01065 |
| 6GQ7 | T857A | DB01076 |
| 6GQ7 | T857A | DB01152 |
| 6GQ7 | T857A | DB01197 |
| 6GQ7 | T857A | DB01229 |
| 6GQ7 | T857A | DB01392 |
| 6GQ7 | T857A | DB01394 |
| 6GQ7 | T857A | DB09054 |
| 6GQ7 | T857A | DB09322 |
| 6GQ7 | T857A | DB11091 |
| 6GQ7 | T857A | glucagon |
| 6GQ7 | T857A | neomycin |
| 6GQ7 | T857A | thyrotropin_releasing_factor |
| 6I83 | R982C | alectinin_hcl |
| 6I83 | R982C | DB00398 |
| 6I83 | R982C | DB00619 |
| 6I83 | R982C | DB00755 |
| 6I83 | R982C | DB01234 |
| 6I83 | R982C | DB01268 |
| 6I83 | R982C | DB01590 |
| 6I83 | R982C | DB05294 |
| 6I83 | R982C | DB08875 |
| 6I83 | R982C | DB08896 |
| 6I83 | R982C | DB08901 |
| 6I83 | R982C | DB09078 |
| 6I83 | R982C | DB09079 |
| 6I83 | R982C | DB11363 |
| 6I83 | R982C | sorafenib_tosylate |

| | | |
|---|---|---|
| 6I83 | R982C | sunitinib_malate |
| 4YHJ | F110V | DB00264 |
| 4YHJ | F110V | DB00661 |
| 4YHJ | F110V | DB00335 |
| 4YHJ | Y292A | DB00264 |
| 4YHJ | Y292A | DB00661 |
| 4YHJ | Y292A | DB00335 |
| 4YHJ | V247I | DB00264 |
| 4YHJ | V247I | DB00661 |
| 4YHJ | V247I | DB00335 |
| 4YHJ | C215I | DB00264 |
| 4YHJ | C215I | DB00661 |
| 4YHJ | C215I | DB00335 |
| 4YHJ | A142V | DB00264 |
| 4YHJ | A142V | DB00661 |
| 4YHJ | A142V | DB00335 |
| 4YHJ | Y53I | DB00264 |
| 4YHJ | Y53I | DB00661 |
| 4YHJ | Y53I | DB00335 |
| 4YHJ | R65L | DB00264 |
| 4YHJ | R65L | DB00661 |
| 4YHJ | R65L | DB00335 |
| 4YHJ | I33L | DB00264 |
| 4YHJ | I33L | DB00661 |
| 4YHJ | I33L | DB00335 |
| 4YHJ | V486A | DB00264 |
| 4YHJ | V486A | DB00661 |
| 4YHJ | V486A | DB00335 |
| 4YHJ | F454A | DB00264 |
| 4YHJ | F454A | DB00661 |
| 4YHJ | F454A | DB00335 |
| 6BDN | S47N | DB00295 |
| 6BFN | L521H | DB00317 |
| 6BFN | L521H | DB00619 |
| 6BFN | L521H | DB00398 |
| 6BFN | F196S | DB00317 |
| 6BFN | F196S | DB00619 |
| 6BFN | F196S | DB00398 |

**Supplemental_Table_S7: Data used for ligand similarity analysis**

| Native | Drug Name | Variation | DrugBank Id | Delta(native-mutant) |
|---|---|---|---|---|
| 6GQ7 | Zinc sulfate | T857A | DB09322 | -9.1 |
| 6GQ7 | Fulvestrant | T857A | DB00947 | -0.3 |
| 5TQY | Pirfenidone | V155A | DB00795 | -0.2 |
| 6GQ7 | Verapamil | T857A | DB00997 | -0.2 |
| 6GQ7 | Sulfasalazine | T857A | DB01076 | -0.2 |

| | | | | |
|---|---|---|---|---|
| 3GC9 | Doxorubicin | R275H | DB04951 | -0.2 |
| 4TNB | Atorvastatin | Q41L | DB00661 | -0.2 |
| 6GQ7 | Mannitol | T857A | DB00742 | -0.2 |
| 6GQ7 | Rizatriptan | T857A | DB00953 | -0.2 |
| 6GQ7 | Mycophenolic acid | T857A | DB01024 | -0.2 |
| 3GC9 | Regorafenib | R275H | DB08896 | -0.1 |
| 4TNB | Trandolapril | Q41L | DB00519 | -0.1 |
| 5TQY | Ascorbic acid | V155A | DB00126 | -0.1 |
| 5TQY | Aminosalicylic acid | V155A | DB00233 | -0.1 |
| 6GQ7 | Lovastatin | T857A | DB00227 | -0.1 |
| 6GQ7 | Liothyronine | T857A | DB00279 | -0.1 |
| 6GQ7 | Progesterone | T857A | DB00396 | -0.1 |
| 6GQ7 | Phenmetrazine | T857A | DB00830 | -0.1 |
| 6GQ7 | Oxazepam | T857A | DB00842 | -0.1 |
| 6GQ7 | Glucagon | T857A | glucagon | -0.1 |
| 6I83 | Octreotide | R982C | DB09078 | 0.1 |
| 6I83 | Lenvatinib | R982C | DB11363 | 0.1 |
| 6GQ7 | Alectinib | T857A | DB00104 | 0.1 |
| 6I83 | Ponatinib | R982C | DB08901 | 0.1 |
| 6I83 | Regorafenib | R982C | DB08896 | 0.7 |
| 6BFN | Sorafenib | F196S | DB00398 | -0.2 |
| 4YHJ | Verapamil | F110V | DB00661 | -0.2 |
| 4YHJ | Atenolol | Y292A | DB00335 | -0.2 |
| 4YHJ | Atenolol | A142V | DB00335 | -0.1 |
| 4YHJ | Atenolol | F454A | DB00335 | -0.1 |
| 4YHJ | Atenolol | V247I | DB00335 | -0.1 |
| 4YHJ | Metoprolol | F454A | DB00264 | -0.1 |
| 4YHJ | Verapamil | F454A | DB00661 | -0.1 |
| 4YHJ | Metoprolol | V247I | DB00264 | -0.1 |
| 4YHJ | Verapamil | V247I | DB00661 | -0.1 |
| 4YHJ | Metoprolol | Y292A | DB00264 | -0.1 |
| 4YHJ | Verapamil | Y53I | DB00661 | -0.1 |
| 4YHJ | Verapamil | C215I | DB00661 | 0.1 |
| 4YHJ | Metoprolol | I33L | DB00264 | 0.1 |
| 4YHJ | Atenolol | R65L | DB00335 | 0.1 |
| 4YHJ | Verapamil | R65L | DB00661 | 0.1 |
| 4YHJ | Metoprolol | Y53I | DB00264 | 0.1 |
| 4YHJ | Atenolol | I33L | DB00335 | 0.3 |
| 6BFN | Gefitinib | L521H | DB00317 | 0.4 |
| 6BFN | Gefitinib | F196S | DB00317 | 0.5 |

**Supplemental_Table_S8**: **Computational prediction of toxicity profiles of drugs using ProTox-II platform**

| DrugBank Ids | Drug Names | Acute Toxicity (mg/kg) | Organ Toxicity | Toxicity endpoints | Adverse outcomes pathways (AOPs) | SIDER ADR links |
|---|---|---|---|---|---|---|

| | | | | | | |
|---|---|---|---|---|---|---|
| DB00947 | Fulvestrant | 2000 | NA | immunotoxic (0.99) | NR_Aromatase (1.00) SR-MMP (0.98) | http://sideeffects.embl.de/drugs/104741/ |
| DB00795 | Sulfasalazine | 1998 | hepatotoxic (0.69) | immunotoxic (0.98) | NR-Aromatase (1.00) NR-ER (0.99) NR-ER-LBD (1.00) | http://sideeffects.embl.de/drugs/5353980/ |
| DB00997 | Doxorubicin | 205 | NA | immunotoxic (0.99) mutagenic (0.98) cytotoxic (0.94) | NR_Aromatase (0.52) SR_p53 (0.52) | http://sideeffects.embl.de/drugs/1690/ |
| DB01076 | Atorvastatin | 5000 | hepatotoxic (0.76) | NA | NR-Aromatase (1.00) | http://sideeffects.embl.de/drugs/2250/ |
| DB04951 | Pirfenidone | 580 | hepatotoxic (0.57) | carcinogenic (0.54) | NA | http://sideeffects.embl.de/drugs/40632/ |
| DB00661 | Verapamil | 108 | NA | immunotoxic (0.97) | NA | http://sideeffects.embl.de/drugs/2520/ |
| DB00742 | Mannitol | 13500 | NA | NA | NA | http://sideeffects.embl.de/drugs/453/ |
| DB00953 | Rizatriptan | 100 | NA | immunotoxic (0.69) | NA | http://sideeffects.embl.de/drugs/5078/ |
| DB01024 | Mycophenolic acid | 800 | hepatotoxic (0.82) | immunotoxic (0.99) cytotoxic (0.77) | SR-MMP (0.79) | http://sideeffects.embl.de/drugs/4272/ |
| DB08896 | Regorafenib | 800 | hepatotoxic (0.82) | immunotoxic (0.99) cytotoxic (0.77) | SR-MMP (0.79) SR-p53(0-57) | http://sideeffects.embl.de/drugs/11167602/ |
| DB00519 | Trandolapril | 1800 | NA | NA | NA | http://sideeffects.embl.de/drugs/5525/ |
| DB00126 | Ascorbic acid | 3367 | NA | NA | NA | http://sideeffects.embl.de/drugs/11020241/ |
| DB00233 | Aminosalicylic acid | 4000 | hepatotoxic (0.82) | NA | NA | http://sideeffects.embl.de/drugs/4649/ |
| DB00227 | Lovastatin | 1000 | NA | carcinogenic (0.80) immunotoxic (0.99) | NR-Aromatase (1.00) SR-MMP (0.88) | http://sideeffects.embl.de/drugs/3962/ |
| DB00279 | Liothyronine | 10000 | NA | NA | NR-AhR (0.94) NR-ER (0.78) NR-PPAR-Gamma (0.94) SR-MMP (0.75) | http://sideeffects.embl.de/drugs/861/ |
| DB00396 | Progesterone | 2450 | NA | carcinogenic (0.56) immunotoxic (0.95) | NR-AR (0.97) NR-LBD (0.97) NR-ER (0.85) SR-ARE (0.86) | http://sideeffects.embl.de/drugs/4920/ |

| | | | | | | |
|---|---|---|---|---|---|---|
| | | | | | SR-p53 (0.80)<br>SR-HSE (0.86) | |
| DB00830 | Phenmetrazine | 125 | NA | NA | NA | |
| DB00842 | Oxazepam | 370 | NA | carcinogenic (0.74) | NR-AR (0.95) | http://sideeffects.embl.de/drugs/4616/ |
| DB09078 | Lenvatinib | 3000 | NA | immunotoxic (0.98) | NA | |
| DB11363 | Alectinib | 2400 | NA | immunotoxic (0.98) | NA | |
| DB00104 | Octreotide | 1000 | NA | immunotoxic (0.74) | NA | http://sideeffects.embl.de/drugs/54373/ |
| DB08901 | Ponatinib | 1190 | NA | immunotoxic (0.96) | NR-Aromatase (1.00)<br>NR-ER (0.99)<br>NR-ER-LBD (1.00) | http://sideeffects.embl.de/drugs/24826799/ |
| DB00317 | Gefitinib | 2935 | hepatotoxic (0.73) | immunotoxic (0.99) | NR-AhR (1.00) | http://sideeffects.embl.de/drugs/123631/ |
| DB00040 | Glucagon | 73 | NA | NA | NA | NA |
| DB00264 | Metoprolol | 1050 | NA | NA | NA | http://sideeffects.embl.de/drugs/4171/ |
| DB00398 | Sorafenib | 800 | hepatotoxic (0.82) | immunotoxic (0.92)<br>cytotoxic (0.77) | NA | http://sideeffects.embl.de/drugs/216239/ |
| DB09322 | Zinc sulfate | 448 | NA | carcinogenic (0.64) | NA | http://sideeffects.embl.de/drugs/24424/ |
| DB00335 | Atenolol | 2000 | NA | NA | NA | http://sideeffects.embl.de/drugs/2249/ |

* aryl hydrogen receptor (AhR), androgen receptor (AR), androgen receptor ligand-binding domain (AR-LBD), aromatase, estrogen receptor alpha (ER), estrogen receptor ligand-binding domain (ER-LBD), and peroxisome proliferator-activated receptor gamma (PPAR-Gamma), Nuclear factor (erythroid-derived 2)-like 2/antioxidant responsive element (ARE), heat shock factor response element (HSE), mitochondrial membrane potential (MMP), phosphoprotein tumor suppressor (p53), NR (Nuclear receptor signalling pathways), SR (Stress response pathways).

*Confidence score: 0.50-0.69 (low), 0.70-0.80 (medium), 0.80-1.00 (high)

**Supplemental_Table _S9: Structural analysis results by DSSP, Naccess, and Dynamut for IndiGen Structure Data.**

| PDB code | Variation | Sec. Str. Assignment (DSSP) | Solv. Acess. (Naccess) | ΔΔG(Dynamut) kcal/mol | ΔΔSVib | Consurf Score |
|---|---|---|---|---|---|---|
| 3D2R | M134V | α-helix | 3 | -0.595 | 0.670 | -0.696 |
| 3GC9 | R275H | Turn | 76.6 | 0.156 | 0.006 | 1.491 |
| 3NRU | I138V | Loop | 13.3 | 0.609 | -0.842 | 1.142 |
| 3NYO | V334M | β-sheet | 28.6 | 0.315 | -0.057 | -0.278 |
| 4EYJ | A300T | α-helix | 40.8 | 0.029 | -0.128 | 1.373 |
| 4TNB | Q41L | 3-10 helix | 47.6 | -0.205 | [illegible]269 | [illegible]5 |
| 4YHJ | F110V | α-helix | 14.5 | -1.024 | 0.844 | 0.512 |
| 4YHJ | Y292A | α-helix | 0 | -0.041 | 1.079 | -1.037 |
| 4YHJ | V247I | β-bridge | 1.8 | 0.379 | -0.181 | -0.851 |
| 4YHJ | C215I | β-sheet | 0 | 0.282 | -0.752 | -0.83 |
| 4YHJ | A142V | α-helix | 18.4 | 0.669 | -0.427 | 1.061 |
| 4YHJ | Y53I | Helix turn | 6.5 | -0.605 | 0.915 | -0.636 |
| 4YHJ | R65L | α-helix | 43.6 | -0.355 | 0.145 | 0.863 |
| 4YHJ | I33L | Loop | 49.8 | 0.640 | -0.133 | -0.156 |
| 4YHJ | V486A | Turn | 76.7 | -0.428 | 0.194 | -0.411 |
| 4YHJ | F454A | α-helix | 14.4 | -2.767 | 1.178 | -0.822 |
| 5TQY | V155A | Loop | 52.1 | -0.622 | 0.073 | 1.941 |
| 6BDN | S47N | Loop | 74.9 | 0.406 | -0.065 | 0.912 |
| 6BFN | L521H | Bend | 17.3 | -0.498 | 0.140 | -0.457 |
| 6BFN | F196S | α-helix | 74.9 | -0.011 | 0.324 | -0.053 |
| 6GQ7 | T857A | α-helix | 70.2 | 0.080 | 0.123 | 0.761 |
| 6I83 | R982C | α-helix | 40.1 | 0.218 | 0.129 | 1.184 |

**Supplemental_Table_S10: PharmaGKB results**

| Native | Variation | Gene | variants | Chemicals | Phenotypes |
|---|---|---|---|---|---|
| 4YHJ | F110V | GRK4 | rs1024323 | metoprolol | Hypertension, Kidney Diseases, Nephrosclerosis |
| 4YHJ | A142V | GRK4 | rs1024323 | metoprolol | Hypertension, Kidney Diseases, Nephrosclerosis |
| 4YHJ | Y292A | GRK4 | rs1801058 | metoprolol | hypertensive nephrosclerosis |
| 4YHJ | V486A | GRK4 | rs1801058 | metoprolol | hypertensive nephrosclerosis |
| 4YHJ | F454A | GRK4 | rs1801058 | metoprolol | hypertensive nephrosclerosis |

**Supplemental_Table _S11: Kinase families associated with 545 kinase coding genes with number of drugs and SNPs observed in each class (Indigen Sequence data).**

| Kinase family | Drug count | SNP count |
|---|---|---|
| ACG | 339 | 2343 |
| ATYPICAL | 148 | 1224 |
| CAMK | 213 | 10581 |
| CK1 | 18 | 275 |
| CMGC | 659 | 1632 |
| STE | 147 | 1734 |
| TK | 1978 | 5073 |
| TKL | 185 | 1193 |
| OTHER | 196 | 4043 |

**Supplemental_Table _S12:** Data used for amino-acid exchange frequency analysis

(Excel sheet- Supplemental_Table _S12.csv)

**Supplemental_Table_S13:** HUMSAVAR variant data used for comparative structural analysis consisting of 217 variants corresponding to 12 genes in IndiGen structure data.
(Excel sheet- Supplemental_Table _S13.xlsx)

**Supplemental_Table _S14:** ADR Profile of Atenolol and Metaprolol

(Excel sheet- Supplemental_Table _S14.xlsx)

**Supplemental_Table _S15:** ADRs and drugs information on the drugs included in our analysis

(Excel sheet- Supplemental_Table _S15.xlsx)
